# Human Stress Response Specificity through Biochemical Resonance Selectivity

**DOI:** 10.1101/2025.03.05.641735

**Authors:** Michael Worcester, Shayan Nejad, Arya Eimagh Naeini, Surya Arian, Devin O’Donnell, Pratyasha Mishra, Siyu Li, Kevin Yang, Aisa Anbir, Matthew Guevara, Nina Y. Yuan, Seán O’Leary, Marcus Kaul, Roya Zandi, Thomas E. Kuhlman

## Abstract

In eukaryotes, the mitogen activated protein kinase (MAPK) cascade, a multilayered interconnected network of enzymes, connects external stimuli to gene regulation, determining cellular fate ^1^. Environmental stress sensed by a cell starts a complex chain of reactions between MAPK enzymes that ultimately activates the master stress response regulator protein p38 MAPK ^2,3^. Thus activated, p38 must then selectively activate targets from a pool of hundreds to initiate appropriate cellular responses ^3^. Mechanisms for how p38 performs this selection remain unclear ^4,5^. Here we show that human p38 target selectivity is based on the same principles as modern electronic telecommunications systems, except using waves of chemicals rather than electromagnetic fields or electric currents. p38 encodes information about stimuli as different frequency oscillations of its activation state, and targets are selected through frequency-dependent resonance of oscillating biochemical reactions between p38 and its targets. We demonstrate this mechanism by activating various genetic responses in human cells by applying only sugar at different frequencies. These results unify observations of oscillating signaling components and altered responses ^6–19^ into a coherent framework to understand and control human gene expression. As failures of this mechanism may contribute to some p38-associated diseases ^2,20–28^, these findings may have implications for pharmaceutical development and therapeutic strategies.

## Main Text

Enzyme specificity is essential for the adaptability and survival of all living organisms. An example relevant to all eukaryotes is the mitogen activated protein kinase (MAPK) cascade, a complex network of dozens of enzymes responsible for sensing environmental stimuli and sending the signal to the genome to alter gene expression. In the case of responding to environmental stresses, the cascade transmits received signals as a current of activating and deactivating phosphorylation reactions between MAPK enzymes, starting from receptors that detect the stress, passed through the MAP3K and MAP2K layers of the cascade, and ultimately received by the master stress response regulator protein p38 MAPK ^2,3^. Once activated, p38 must then selectively phosphorylate and activate the correct substrates from a pool of hundreds of different possible protein targets to initiate gene expression responses appropriate to the received stress ^3^. Selection of incorrect responses can be catastrophic, as inappropriate p38 activity is implicated in many devastating human diseases, including Alzheimer’s disease ^29^, amyotrophic lateral sclerosis (ALS) ^30^, neuroHIV ^31^, and cancer ^20^, among many others ^2,32^.

How can p38, which primarily exists only in either an active phosphorylated state or an inactive dephosphorylated state ^2^, encode enough information to provide specificity in response selection? In some cases, specificity in p38-target interactions is aided by allosteric control ^33^ or regulation of p38 and substrate availability, localization, or scaffolding ^4,34,35^. However, because p38 regulates hundreds of different targets, it is difficult to imagine how to generalize such approaches to form a comprehensive model of p38 selectivity due to the numbers of additional allosteric and localization effectors that would be required but remain unidentified.

Extracting order from this complex network is further complicated by dynamic temporal patterns and oscillations in number and activation state that have been observed in thousands of proteins within and associated with the MAPK cascade ^36^. These include the master regulator ERK1/2 ^6,7^, the MAP2Ks MKK3 and MKK6 ^8–11^, and the transcription factors (TFs) NF-κB ^12,13^ and p53 ^14,15^, among many others ^16^. Because the output of a particular signaling pathway relies on the balance between activation and deactivation by kinase and phosphatase enzymes, the temporal behavior of pathway components has been proposed to contribute to signaling specificity ^12,13,17^. However, specific mechanisms for how such temporal dynamics are interpreted or employed by signaling pathways remain largely obscure.

### VPC (Venus-p38-Cerulean) directly reports single cell p38 activation state by FRET

Existing tools for studying information flow through the MAPK cascade and p38, such as kinase translocation reporters (KTRs)^18^ and Förster resonance energy transfer (FRET) based kinase substrates, rely upon indirect measurements. An example is the FRET-based p38 substrate PerKy-38 ^19^, which is based upon the p38 docking domain of the TF protein MEF-2A and changes its FRET state upon phosphorylation by p38. However, while such reporters are phosphorylated by p38, they are dephosphorylated independently by other phosphatases ^37,38^. Hence PerKy-38 and KTRs report on the activation state dynamics of p38 substrates rather than those of p38 itself.

To overcome this limitation, we developed VPC (Venus-p38-Cerulean), which directly reports p38 activation state by FRET, and we validated that the phosphorylation and activity dynamics of VPC are representative of native p38 on the time scales measured here. VPC is a fusion of the N and C terminal ends of p38α, the most ubiquitous p38 isoform ^39^, to the fluorescent reporters mVenus ^40^ and mCerulean3 ^41^. When human HEK-293T cells stably transfected to express low levels of VPC (henceforth VPC293) are stimulated, intracellular VPC displays an increased FRET signal (**Fig. 1a,b)**. VPC phosphorylation state dynamics measured by western blot and averaged FRET recapitulate those of p38 in untransfected HEK-293T cells (**Fig. 1c, Extended Data Fig. 1a**). Furthermore, the FRET signal of VPC purified from unstimulated cells increases when phosphorylated *in vitro* by the MAP2K p38 kinase MKK6 (**Fig. 1d, Extended Data Fig. 1b,c**). Finally, the FRET state of stimulated VPC293 cells changes dynamically in time (**Fig. 1e**) and in amplitude (**Fig. 1f**) in response to osmotic shocks of various strengths.

**Fig. 1.**
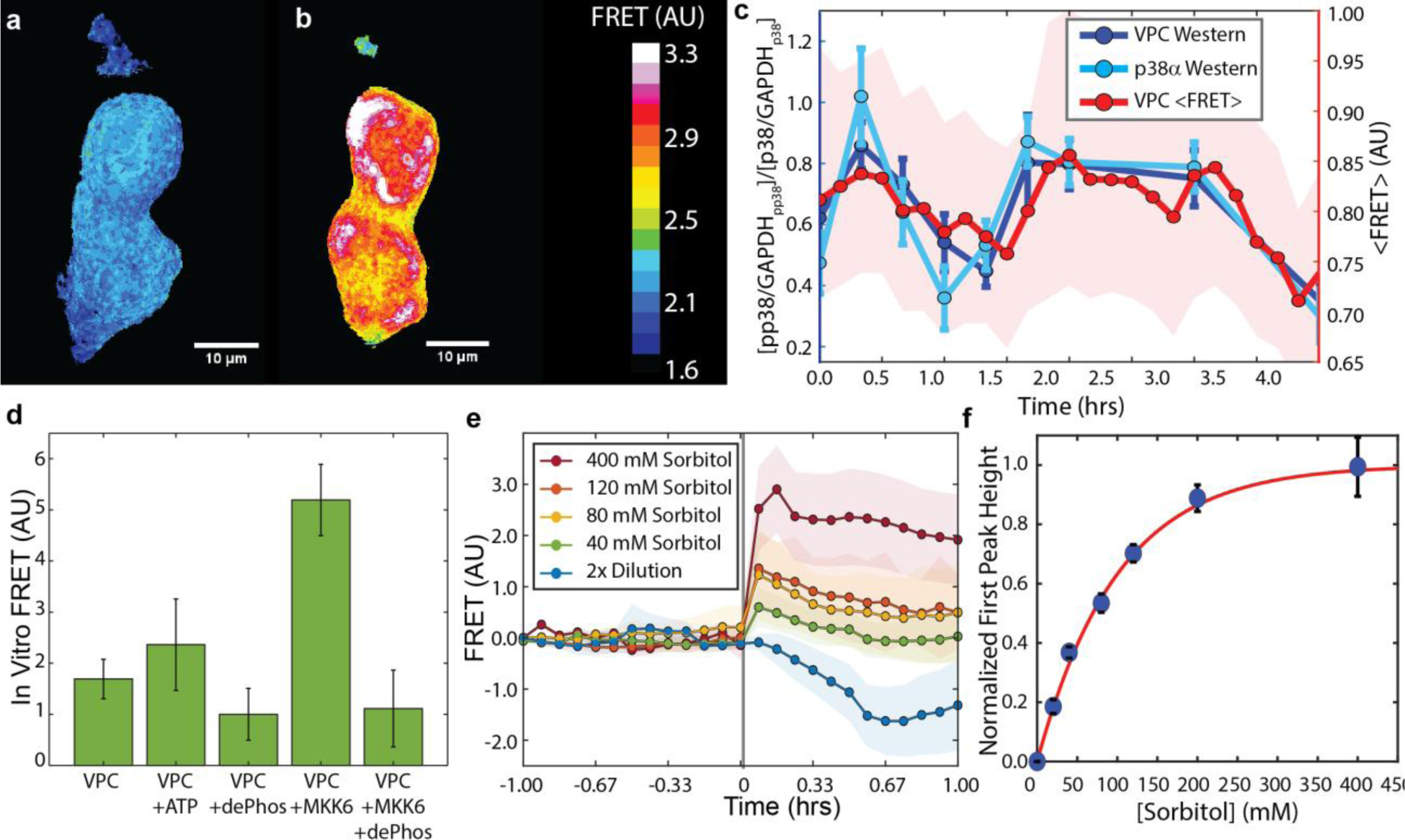
VPC, a direct FRET-based reporter of p38 activation state. (**a**) FRET state of two HEK-293T cells expressing VPC (VPC293). (**b**) The same two cells, immediately after hyperosmotic shock induced by 120 mM sorbitol. (**c**) VPC (dark blue, from VPC293 cells) and p38α (light blue, from untransfected HEK-293T cells) phosphorylation dynamics measured using Western blot and average FRET (red, *N* ≈ 1,500 cells). Shaded region is the 95% confidence bound for FRET data, while error bars are standard deviation of three sets of western blot biological replicates. See **Extended Data Fig. 1** for example western blots. (**d**) *In vitro* FRET of purified VPC protein; as purified (VPC), +0.25 mM ATP-Mg (VPC +ATP), VPC +0.25 mM ATP-Mg +100 U/ml Antarctic Phosphatase (VPC +dePhos), VPC +0.25 mM ATP-Mg +200 nM MKK6 kinase (VPC +MKK6), and VPC +0.25 mM ATP-Mg +200 nM MKK6 +100 U/ml Antarctic Phosphatase. Values are means of 15 measurements, error bars are SEM. (**e**) Dynamic averaged FRET response of VPC293 cells upon hyperosmotic (Green: 40 mM, yellow: 80 mM, light red: 120 mM sorbitol, dark red: 40 mM) or hypoosmotic (Blue: 2x dilution with ultrapure H_2_O) shock. Shock was induced at *t* = 0, indicated by the vertical gray line. Points: average of three biological replicates, shaded region indicates 95% confidence bounds. (**f**) Normalized height of the first peak in the VPC FRET signal upon induction of hyperosmotic shock via the addition of sorbitol, as in (e). Blue points: mean of three biological replicates, error bars: standard deviation. Red line is best fit to the function 1 − *e*^−*b*∗[Sorbitol]^, with the best fit value of *b* = 0.01 ± 0.002 mM^-1^. Hyperosmotic shock with 400 mM sorbitol results in cell death.

### Measuring information flow as enzyme activity oscillations in the MAPK cascade

By measuring the FRET responses of both p38 and the TF-based p38 substrate PerKy-38 in response to stimuli, we directly observe how information is passed between layers of the MAPK cascade at the single cell level. We find that, when stimulated, the activation states of both p38 and PerKy-38 in individual cells oscillate in time (**Fig. 2a-c**), corresponding to temporal waves in numbers of active molecules.

**Fig. 2.**
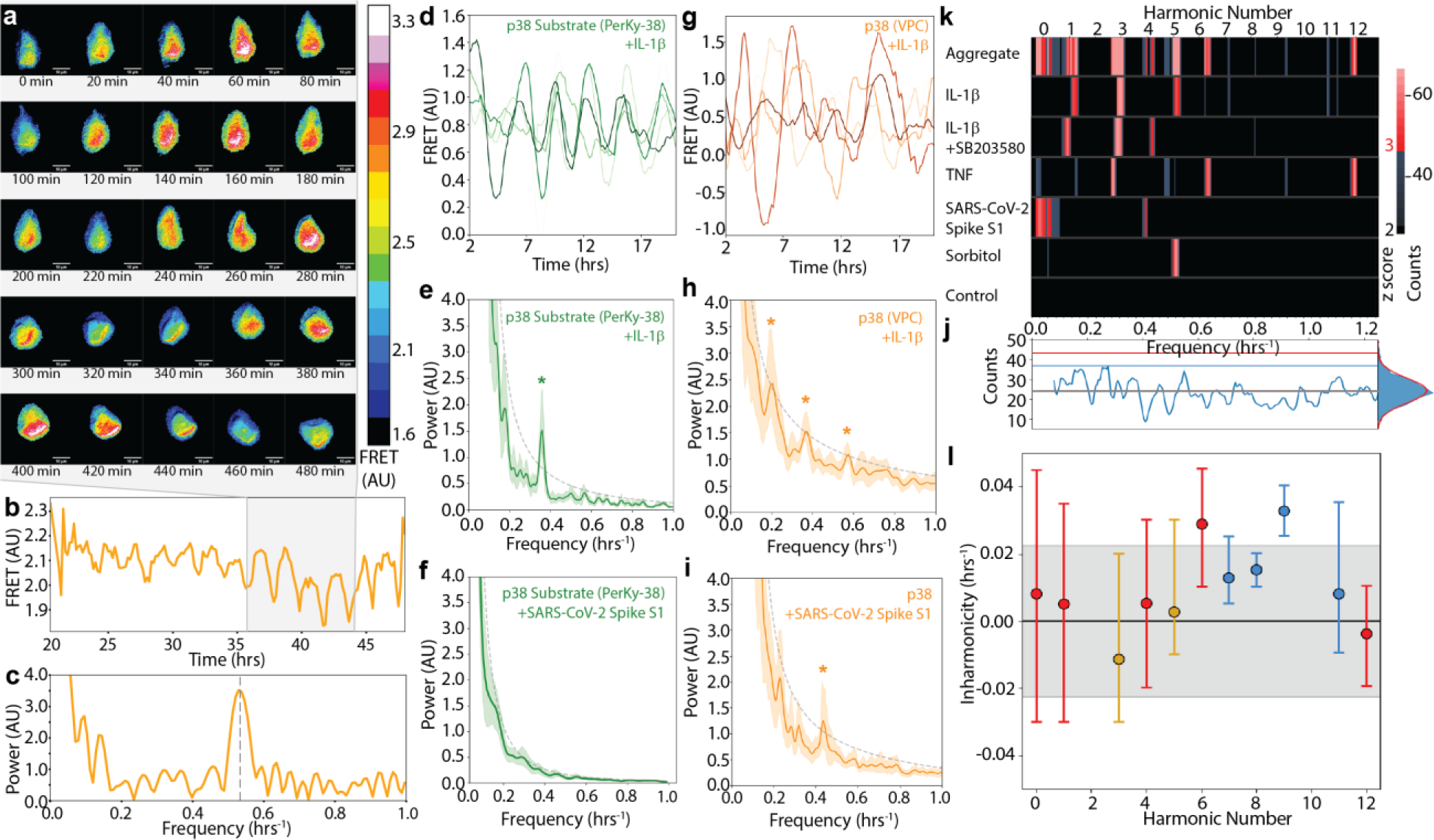
p38 and PerKy-38 exhibit phosphorylation state oscillations with stimulus-specific frequencies. (**a**) Oscillating p38 activation state of a VPC293 cell stimulated by hyperosmotic shock. The first image (t = 0) is ∼36 hours after addition of 120 mM sorbitol. Scale bar: 10 μm. (**b**) FRET time trace and (**c**) Fourier spectrum of the cell shown in (a). Shaded region indicates time period of images in (a); dashed line in (c) indicates *f* = 0.54 hr^-1^. (**d**) Four FRET time traces of PerKy293 cells +300 ng/mL IL-1β. (**e**) Average Fourier power spectrum of *N* = 475 PerKy293 cells +300 ng/mL IL-1β. Shaded region: 95% confidence bounds; dashed gray line: 99% confidence interval for peak magnitudes. Asterisks: statistically significant peaks. (**f**) Average spectrum of *N* = 750 PerKy293 cells +2 nM SARS-CoV-2 Spike S1. (**g**) Four FRET time traces of VPC293 cells +300 ng/mL IL-1β. (**h**) Average spectrum of *N* = 495 VPC293 cells +300 ng/mL IL-1β. (**i**) Average spectrum of *N* = 525 VPC293 cells +2 nM SARS-CoV-2 Spike S1 protein. (**j**) Statistics of local maxima in Fourier spectra of unstimulated VPC293 cells (see **SM II**). Frequency integration yields a normal distribution (right) used to quantify *z* score; *z* = number of standard deviations from mean. Gray: mean; blue: *z* = 2; red: *z* = 3. (**k**) Oscillatory frequencies for different stimuli. Top: aggregated spectrum. Expected frequencies of the *n*^th^ harmonic [*f_n_* = (*n* + 1)*f*_0_, *n* = 0, 1, 2, …, and *f*_0_ = 0.09 hr^-1^] indicated. Blue: 2 ≤ *z* < 3 and red: *z* ≥ 3. (**l**) Inharmonicity of identified frequencies. Points: band center of mass, error bars: band width at *z* score boundary. Blue: 2 ≤ *z* < 3; red: *z* ≥ 3; yellow: *z* > 10. Gray shaded region: interval of ± *f*_0_/4 (see **SM III**).

### PerKy-38 activity oscillations at a single frequency result from only some stimuli

Example oscillatory time traces from HEK-293T cells stably transfected to express PerKy-38 (henceforth PerKy293) when stimulated with the pro-inflammatory cytokine interleukin-1β (IL-1β) are shown in **Fig. 2d**. The response is eliminated upon treatment of the cells with SB203580 (**Extended Data Fig. 2**), a p38-specific inhibitor that does not interfere with phosphorylation of p38, but which inhibits phosphorylation of downstream targets by p38 ^42^. We performed Fourier analysis on the FRET time traces of 750 cells recorded over 48 hours [see **Supplementary Methods (SM) I**]. These cells are highly motile (**Supplementary Movie 1**), introducing noise into cell tracking and spurious signals in individual cell spectra; we therefore averaged together individual cell spectra (**Extended Data Fig. 3**), leaving detailed characterization of noise and possible response heterogeneity for future work. The average response of the SB203580-treated PerKy293 control exhibits a smooth spectrum of noise, and quantification of the statistics of fluctuations from average noise behavior allows us to determine statistical significance of spectral peaks (see **SM II**, **Extended Data Fig. 2**). For PerKy293 cells stimulated with IL-1β, we find the only statistically significant peak at a frequency of *f* = 0.36 ± 0.02 hr^-1^ (*p* = 2.5 x 10^-13^), corresponding to a period of ∼2.8 hours (**Fig. 2e**). Similarly, we observe a single statistically significant peak at *f* = 0.36 ± 0.03 hr^-1^ when cells are stimulated with the multipurpose cytokine Tumor Necrosis Factor (TNF, *p* = 2.9 x 10^-2^, **Extended Data Fig. 4a,b**). Conversely, we observe no PerKy-38 activity or oscillations when cells are exposed to SARS-CoV-2 Spike S1 protein (**Fig. 2F**) or when osmotically shocked by addition of 120 mM sorbitol to the medium (**Extended Data Fig. 4c,d**).

### p38 activity oscillates with stimulus-specific harmonic frequencies

The p38 activation state in VPC293 cells also shows an oscillatory response when stimulated (**Fig. 2g**). However, unlike PerKy-38, p38 oscillations occur with all stimuli used, but at different frequencies depending on the stimulus. As a control, cells expressing both singly labeled Venus-p38 and p38-mCerulean3 (“VPPC293”) exhibit a spectrum of only noise (**Extended Data Fig. 5, SM II**). Similarly to PerKy-38, the average spectrum from VPC293 cells stimulated with IL-1β displays a statistically significant peak at 0.36 ± 0.02 hr^-1^ (*p* = 4.6 x 10^-3^), but additionally shows other significant peaks at 0.18 ± 0.02 hr^-1^ (*p* = 2.9 x 10^-3^) and 0.54 ± 0.02 hr^-1^ (*p* = 3.1 x 10^-3^) (**Fig. 2h**). When exposed to SARS-CoV-2 Spike S1, the VPC spectrum displays a statistically significant peak at 0.45 ± 0.01 hr^-1^ (*p* = 5.7 x 10^-4^; **Fig. 2i**). Hyperosmotic shock with 120 mM sorbitol results in a peak at a frequency of 0.54 ± 0.02 hr^-1^ (*p* = 2.4 x 10^-4^). Combining these individual spectra (**Fig. 2j,k**) into an aggregate spectrum (**Fig. 2k**, top) shows that every detected frequency from all stimuli tested here resemble evenly spaced harmonics, *f*_*n*_ = (*n* + 1)*f*_0_, with *n* an integer. Quantitative analysis extracts a best-fit fundamental frequency of *f*_0_ = 0.090 ± 0.002 hr^-1^ (see **SM III** and **Extended Data Fig. 6a,b**) and shows that p38 frequencies are distributed closer to harmonics than would be expected at random (*p* = 0.004; see **Fig. 2l**, **Extended Data Fig. 6c**, **SM III**). Accordingly, for the tested stimuli we only observe a response in PerKy293 cells with the cytokines IL-1β or TNF, which activate p38’s 3^rd^ harmonic at *f*_3_ = 0.36 hr^-1^, but observe no response otherwise (**Fig. 2d-f**, **Extended Data Fig. 4**).

### p38 activates PerKy-38 through biochemical resonance

The spectra of p38 and PerKy-38 exhibit exactly the relationship expected between two oscillators coupled by the phenomenon referred to in physics as resonance ^43^, where, similar to pushing a swing, a target that oscillates at a preferred natural frequency (PerKy-38) undergoes large amplitude oscillations only when pumped by a driving force (p38) oscillating at the same frequency [see **Supplementary Discussion (SD) IV, SD V**]. We propose to call this phenomenon biochemical resonance, or bioresonance, to distinguish it from unrelated uses of the term resonance in chemistry and biochemistry.

The relationship between p38 and PerKy-38 is quantified by the transfer function, *Y*(ω), which describes how the PerKy-38 output spectrum, *x̃*(ω), arises from input of the p38 spectrum, *z̃*(ω), such that *x̃*(ω) = *Y*(ω)*z̃*(ω) (see **SD VI**). For linear oscillators coupled by resonance, *Y*(ω) is a known complex function that concisely quantifies all aspects of the interaction, including phase relationships, transfer of energy between the oscillators, and amplitude of the response (see **SD VI**). We find the p38/PerKy-38 transfer function *Y*(ω) is precisely predicted at all frequencies as a resonant interaction between coupled oscillating biochemical reactions (**Fig. 3a-c**; see **SD VI**). Fitting the expected form of *Y*(ω) to the measured response extracts parameters of the resonant interaction, showing that the damping rate from spontaneous dephosphorylation is low [*γ* = 0.00423 ± 0.00009 (SEM) hr^-1^; see **SD IV-V**] and that the Quality factor, *Q*, which measures the magnitude and specificity of resonant amplification (see **SD IV**), is high for such low frequency oscillations, *Q* = 535 ± 11.

**Fig. 3.**
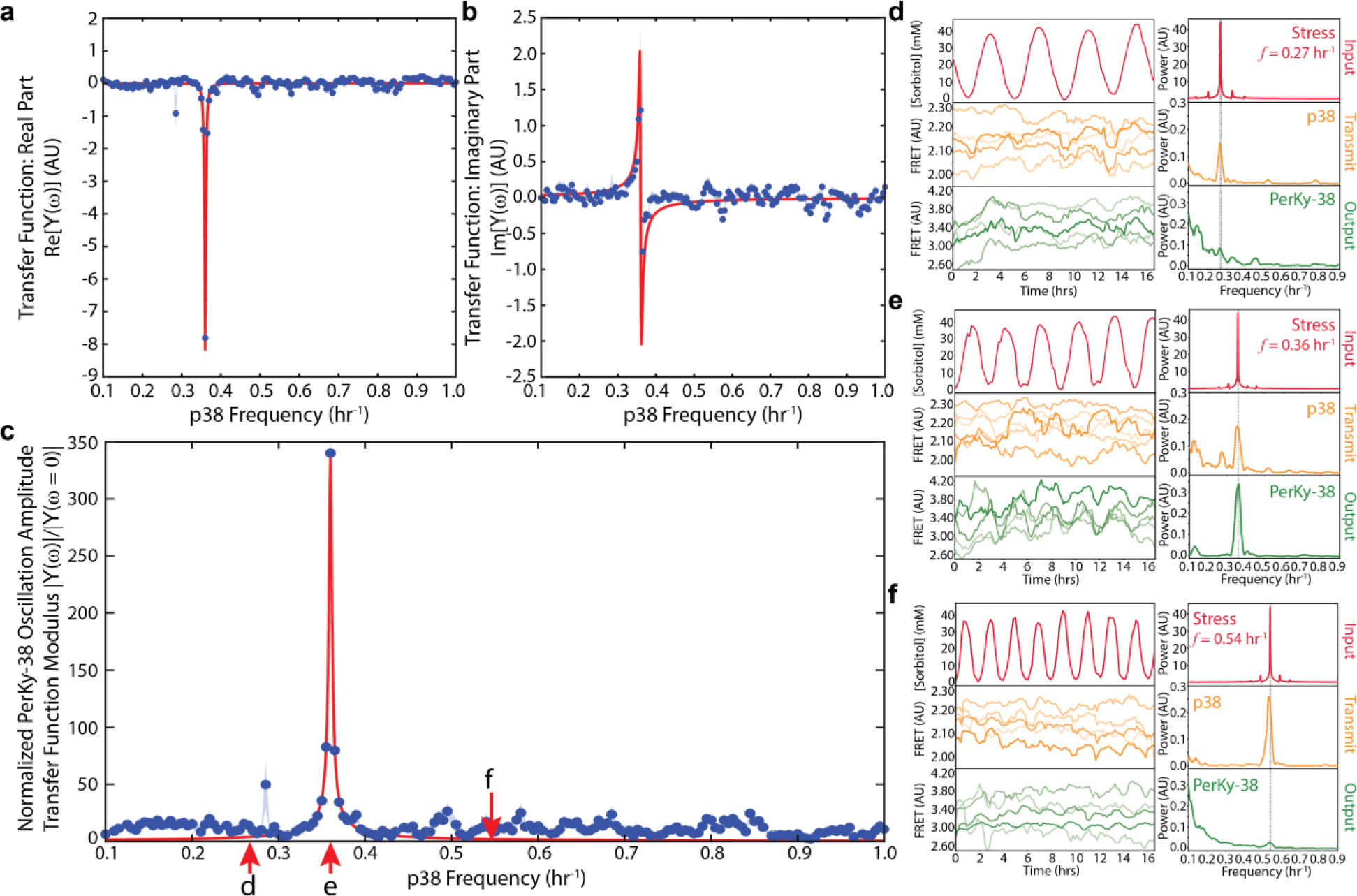
p38 drives target activation through forced resonant oscillations. Real (**a**) and Imaginary (**b**) parts of the transfer function, *Y*(ω), between p38 [*z̃*(ω)] and PerKy-38 [*x̃*(ω)], *Y*(ω) = *x̃*(ω)/*z̃*(ω), see **SD VI**. Blue points: data, *N* = 1,092 ratios of Fourier spectra from VPC293 and PerKy293 cells; blue shaded region: 95% confidence region; red lines: fits to expected form of the transfer function of a driven simple harmonic oscillator; see **SD VI**, eqs. S6.2.3, S6.8.2. (**c**) Normalized modulus of the transfer function between p38 and PerKy-38, showing PerKy-38 amplitude response as a function of p38 driving frequency, see **SD IV**, eq. S4.5, and **SD VI**, eq. S6.10.2. Fits yield damping parameter *γ* = 0.00423 ± 0.00009 (SEM) hr^-1^, quality factor, *Q* ≡ ω_0_/γ = 535 ± 11. (**d-f**) Driving the p38 MAPK system with periodic hyperosmotic shock. Time series (left) and Fourier spectra (right) of the input sorbitol flow (red, top, measured by rhodamine B fluorescence), p38 response (orange, middle, VPC FRET), and PerKy-38 output (green, bottom, PerKy-38 FRET) when cells are subjected to periodic hyperosmotic stress by oscillatory flow of sorbitol between 0 mM and 40 mM at a frequency of (**d**) 0.27 hr^-1^, (**e**) 0.36 hr^-1^, and (**f**) 0.54 hr^-1^. The corresponding driving frequencies relative to the modulus of the transfer function are indicated by red arrows in (c).

### Forcing p38 oscillations of a specific frequency

We generalized our observations of the frequency-based specificity of the p38/PerKy-38 interaction to hypothesize that different frequencies in the p38 oscillatory spectrum resonate with other natively occurring targets with corresponding natural frequencies to selectively drive responses to various stimuli. This suggests it should be possible to elicit different responses by stimulating cells to force p38 oscillations of a specific frequency, regardless of the precise physical or chemical nature of the stimulus.

To test this hypothesis, we took advantage of the linear relationship between low sorbitol input concentrations and the resulting p38 response output amplitude (**Fig. 1f**) by using a pump system designed to deliver a low-concentration sorbitol solution inducing weak sinusoidally varying osmotic shock with a predetermined frequency (**Extended Data Fig. 7**). When cells are stimulated by sorbitol concentration oscillating between 0-40 mM at off-resonant frequencies, p38 oscillates at that frequency in VPC293 cells, while there is no or very weak response in PerKy-38 (**Figs. 3d,f**). Conversely, when cells are driven at PerKy-38’s resonant frequency of *f* = 0.36 hr^-1^, both the VPC and PerKy-38 spectra exhibit statistically significant peaks at that frequency (**Fig. 3e)**, consistent with PerKy-38 activation due to bioresonance.

### Bioresonant stimulation radically alters overall gene expression

RNA sequencing demonstrates that periodically driven cells exhibit radically altered gene expression profiles when p38 oscillations are forced at different frequencies. We collected and sequenced RNA from HEK-293T cells where for 48 hours the sorbitol concentration oscillated between 0 - 40 mM at all p38 harmonic frequencies from 0.09 hr^-1^ to 0.90 hr^-1^. We additionally sequenced RNA from untreated samples as a negative control, samples treated continuously with sorbitol at 20 mM or 40 mM for 48 hours (the mean and maximum sorbitol concentrations of the oscillations, respectively), and samples driven periodically at 0.36 hr^-1^ also treated with the p38-specific inhibitor SB203580. Cluster and principal component analysis of genes differentially expressed relative to the negative controls shows that the negative controls and continuously treated samples exhibit similar expression profiles (**Fig. 4a,b**, **Extended Data Fig. 8**). Conversely, expression profiles of periodically stimulated cells differ dramatically from those treated with continuous stimulation, and from each other depending on the driving frequency, with unique blocks of genes differentially regulated at each driving frequency (**Fig. 4a,b**, **Extended Data Fig. 8**). In particular, the samples driven at 0.36 hr^-1^ and 0.54 hr^-1^, the two most statistically significant frequencies in this HEK-293T cell line’s Fourier spectra (**Fig. 2k-l**), exhibit unique blocks of ∼500 differentially expressed genes (**Fig. 4c**). The samples driven periodically at 0.36 hr^-1^ and treated with SB203580 show generally lower expression of relevant genes compared to all other samples, verifying the bioresonance effect on overall gene expression measured here is mediated by p38.

**Fig. 4.**
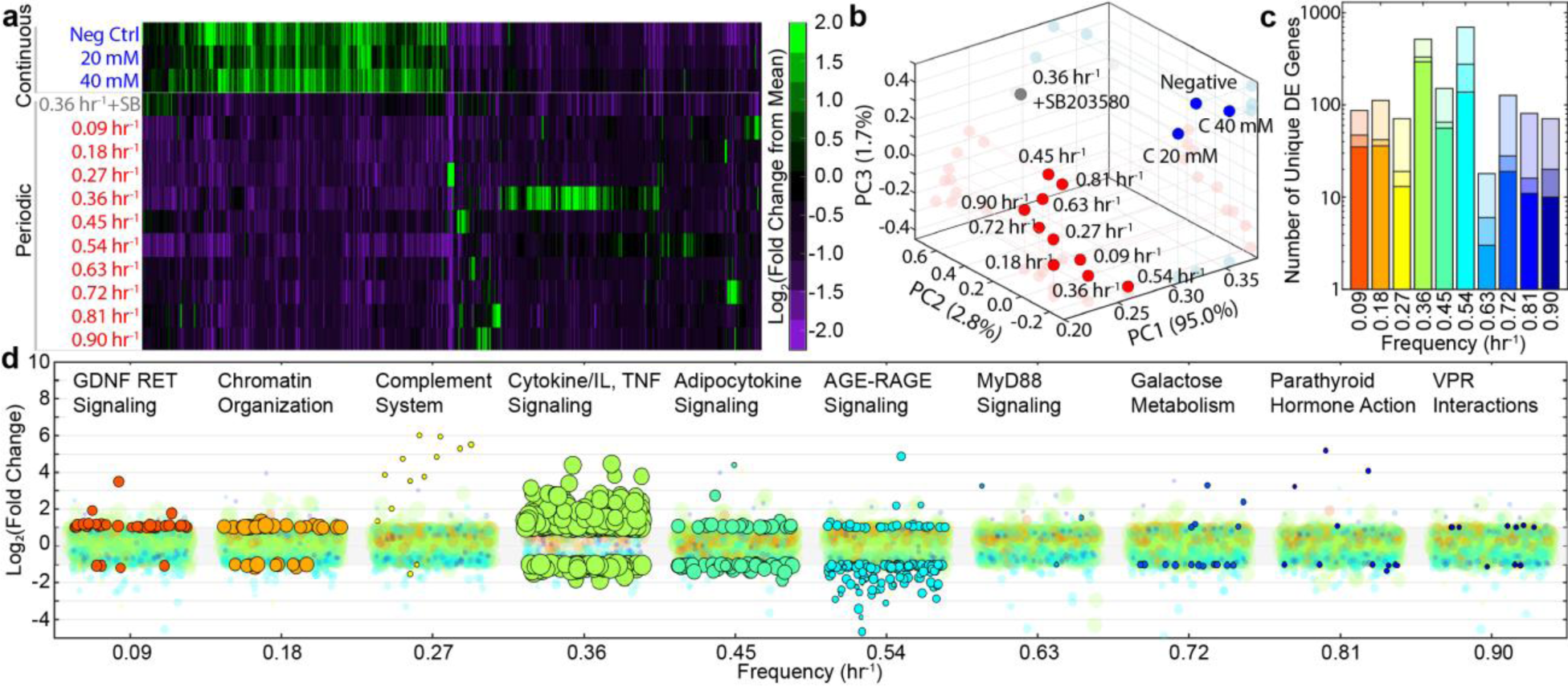
Activation of specific genetic responses through dynamic bioresonant stimulation. (**a**) Clustered heat map of 1,930 identified statistically significantly differentially expressed [DE, *p_adj_* ≤ 0.05, |log_2_(fold change)| ≥ 1] genes uniquely affected when driven at each indicated frequency, as in **Fig. 3d-f**. Color scale is normalized to the mean across all samples. (**b**) Principal Component (PC) analysis of all 58,735 genes identified through RNAseq. See also **Extended Data** Fig. 8. (**c**) Number of genes uniquely DE at each driving frequency. Lightest shade colors: all unique DE genes (*p_adj_* ≤ 0.05, |log_2_(fold change)| ≥ 1); Intermediate shade: highly statistically significant unique DE genes (*p_adj_* ≤ 0.005-0.01, |log_2_(fold change)| ≥ 1); Darkest shade: highly statistically significant unique DE genes intersecting at least one enrichment term. (**d**) Expression level as a function of driving frequency of highly statistically significant unique DE genes intersecting at least one annotated enrichment term. Color scheme corresponds to (c). Points not statistically significant are transparent, while statistically significant points are opaque with black outline. Point size is proportional to number of enrichment terms intersected by that gene. The horizontal position with which each point is plotted within a given frequency band is random. Example enriched pathways at each driving frequency are provided above each frequency band; complete enrichment results are available in **Supplementary Table 2**.

### Distinct genetic programs are activated by different p38 oscillatory frequencies

Functional enrichment analysis demonstrates that forcing p38 oscillations at different frequencies activates distinct genetic programs. From those genes identified as uniquely significantly differentially expressed at each frequency (**Supplementary Table 1**), we performed enrichment analysis on the most highly statistically significant genes, *i.e.*, those with *p*_adj_ ≤ 0.005 for all frequencies except 0.63 hr^-1^, where we used *p*_adj_ ≤ 0.01 to maintain enough genes for analysis. We find that each driving frequency activates unique genetic programs with known associations to p38 in a wide variety of contexts (**Fig. 4d**, **Supplementary Table 2**). Of particular note, differentially expressed genes driven at 0.36 hr^-1^, a frequency that we found appears naturally in response to the cytokines IL-1β and TNF (**Fig. 2k**) and results in activation of PerKy-38, are associated with MAPK responses to cytokines generally and interleukin and TNF specifically, as well as inflammation, regulation of cell death genes, and other processes known to be regulated by p38 (**Fig. 4d** and **Supplementary Table 2**). Moreover, the TF MEF-2A, which forms the molecular basis of PerKy-38, is found to be statistically significantly associated with enriched pathways only at driving frequency of 0.36 hr^-1^ (*p*_adj_ = 5.0 x 10^-4^), consistent with direct FRET-based results. Intriguingly, pathways identified at several frequencies are associated with a variety of diseases where inappropriate p38 activity is implicated ^2,20,29–32^, including cancer (0.09 hr^-1^, 0.36 hr^-1^, 0.45 hr^-1^, and 0.54 hr^-1^), HIV Viral Protein R (VPR) interactions ^31,44^ (0.90 hr^-1^, *p*_adj_ = 8.9 x 10^-3^), and multiple neurodegenerative disorders including neuroinflammation (0.36 hr^-1^, *p*_adj_ = 2.8 x 10^-4^), ALS (0.36 hr^-1^, *p*_adj_ = 4.5 x 10^-3^), and Alzheimer’s disease (0.36 hr^-1^ and 0.45 hr^-1^).

## Discussion

Enzyme specificity is canonically thought of in terms of lock and key mechanisms, where an enzyme’s shape selectively conforms only to that of a specific substrate. That shape can be modified through allosteric or covalent modifications, or specific interactions can be encouraged by colocalizing or scaffolding the enzyme and a subset of targets. Instead, with biochemical resonance a single enzyme can be used to specifically control a broad range of targets simply by altering the frequency with which the amount of active enzyme oscillates. Only those substrates that naturally oscillate due to their other interactions in the cell at the same frequency as the driving enzyme’s oscillations will be amplified through bioresonance. Exploration of mechanisms generating MAPK oscillations is beyond the scope of the present work, but biochemical interactions leading to such oscillations in p38 and other systems are well characterized ^14,15,19,45–48^ (see **SD V**). As we show here, bioresonance is sufficient to provide selectivity and can specifically control gene expression through standard models of transcriptional regulation by protein transcription factors (see **SD VII**, **Extended Data Fig. 9**). This does not preclude additional allosteric, scaffolding, or localization mechanisms to enhance specificity. Furthermore, these results suggest that the frequencies with which thousands of proteins have been observed to oscillate ^6–19^ may determine which interactions are enhanced or suppressed under various conditions. We therefore hypothesize that, in addition to the interactome map of protein-protein interactions, there exists an “oscillatiome,” or a catalog of frequencies with which components oscillate that determines which interactions are differentially affected by bioresonance.

Our mathematical description of bioresonance (**Fig. 5a-e, SD V-VIII**) matches that describing the behavior of electronic alternating current (AC) circuits employed in telecommunications (**Fig. 5f, SD IX-XI**), where resonance selectivity is frequently used as a control scheme. For example, in a simple radio receiver, a desired radio signal is selected from the multitude of transmitted signals by tuning the oscillatory frequency of an AC circuit within the receiver to resonate with the desired signal’s electromagnetic wave oscillations. In the language of AC circuits, tuning the circuit to be in resonance with the radio wave minimizes the frequency dependent impedance, *Z*(ω) ≡ *Y*^−1^(ω) (see **SD IX**), resulting in large amplitude current oscillations that are decoded as audio information. Our results show that the p38 MAPK system functions through a biochemical manifestation of this same resonance selectivity mechanism. As many MAPK components oscillate similarly to that described here for p38 and its substrates ^6–19^, we hypothesize that the MAPK cascade as a whole functions as a complex electronic AC circuit, amplifying and distributing a literal electrical oscillating current of negatively charged phosphate residues to TFs responsible for controlling gene expression (see **SD IX-X** and **Fig. 5g**). The magnitude of the phosphate current received by each TF is determined by the net impedance to the flow of phosphate current, or the product of transfer functions, through each element along the path (**Fig. 5g**; see **SD X**). This, in turn, determines the degree to which regulated genes are differentially expressed.

**Fig. 5.**
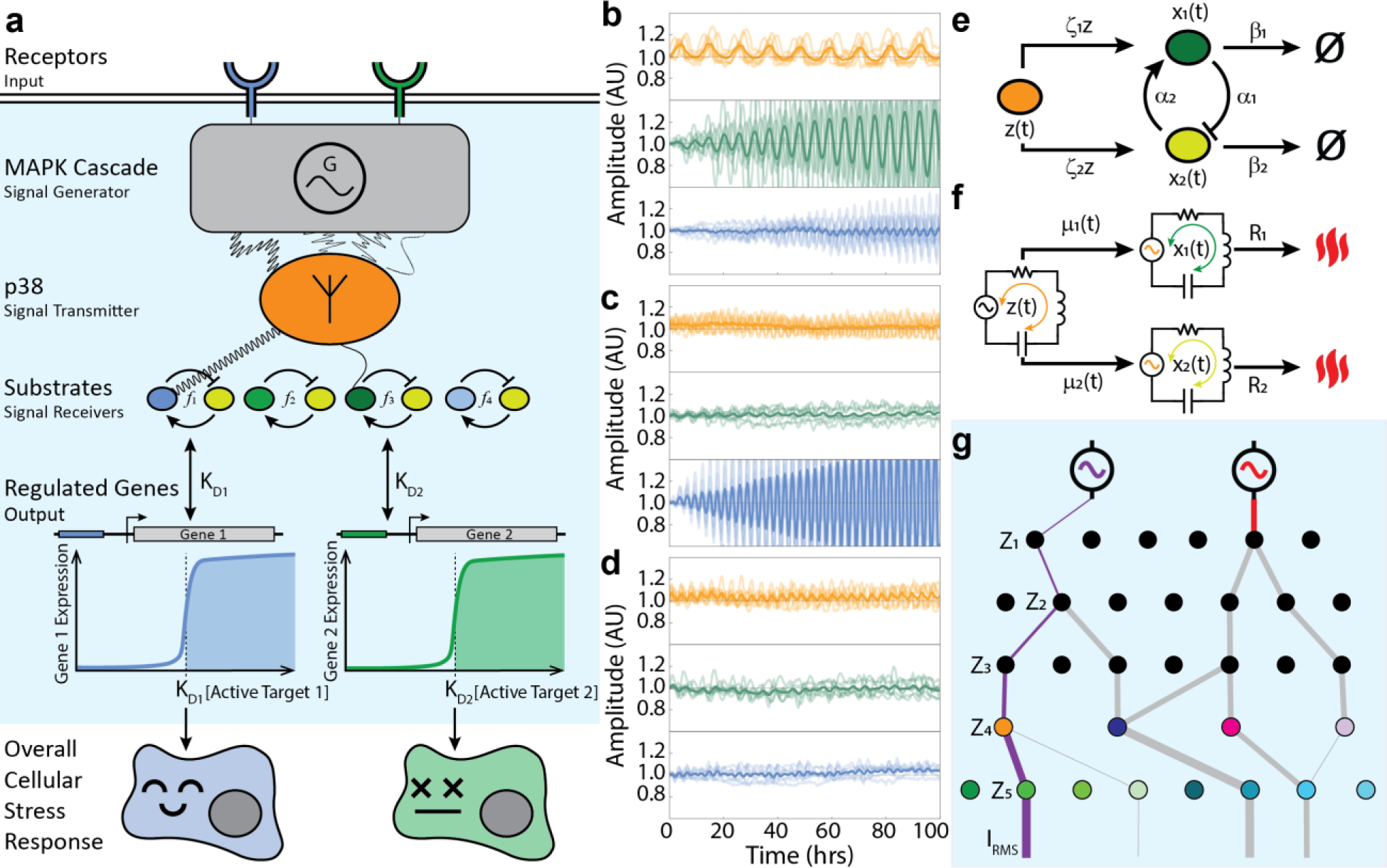
An Integrated Conceptual Framework Describing MAPK Bioresonance and Output. (**a**) Stimulated receptors activate the MAPK cascade, which we treat as a “black box” signal generator (generator symbol) driving p38 oscillations (orange oval) at different frequencies. p38 functions as a transmitter (antenna symbol) that resonantly drives oscillations of downstream targets (green, blue, and yellow ovals), which we model as two-component biological oscillators with natural frequencies *f_i_*. Resonantly activated substrates then differentially regulate their controlled genes. The combined changes in gene expression lead to the overall correct (blue) or incorrect (green) cellular stress response. (**b - d**) Simulated time traces of p38 (orange) resonantly interacting with two downstream targets, Substrate A and Substrate B, with natural frequencies *f*_A_ = 0.18 hr^-1^ (green) and *f*_B_ = 0.36 hr^-1^ (blue). Shown are examples with: (**b**) p38 driven at *f* = 0.09 hr^-1^, which generates harmonics on-resonance with Substrate A; (**c**): p38 driven at *f* = 0.36 hr^-1^, on-resonance with Substrate B; (**d**) p38 driven at *f* = 0.27 hr^-1^, off-resonance with both substrates. Transparent lines: simulations of individual cells; solid lines: mean of seven simulations. See **SD VIII**. (**e**) A quantitative model of p38 [*z*(*t*), orange] driving oscillations in two interacting TFs [*x*_*i*_(*t*), green and yellow]. TFs are phosphorylated with rate constants ς_*i*_; TFs react with each other with rate constants α_*i*_; TFs spontaneously dephosphorylate (null symbol) with rates β_*i*_. See **SD V**. (**f**) Equivalent representation of (e) in terms of AC circuit elements, oscillating phosphate current in p38 (orange) inducing potentials, μ_*i*_(*t*), in downstream TFs, with spontaneous dephosphorylation analogous to resistive heat losses. See **SD V** and **SD IX – X**. (**g**) Representation of the MAPK cascade as a phosphate current amplifier and distribution network. Phosphate currents (I_RMS_, lines) are amplified or suppressed along paths through the cascade based upon the net frequency-dependent impedance, *Z*, of each MAPK element (circles) along each path.

This correspondence to telecommunications strategies suggests that other electronic information encoding and transmission approaches could be relevant to the cell. For example, p38’s use of harmonic frequencies to transmit information may be significant, as harmonic frequencies are orthogonal and result in minimal crosstalk between channels ^49^ (see **SD XI**). The same modulation scheme, referred to as frequency shift keying with orthogonal frequency-division multiplexing (FSK-OFDM) (see **SD XI**), is used in Wi-Fi, television and audio broadcasting, and wireless cellular communications for precisely this reason ^50^.

Finally, this quantitative understanding of the bioresonant circuit properties of the MAPK cascade allows us the mechanistic insight to control human genetic responses using nothing more than low concentrations of the innocuous sugar alcohol sorbitol applied at different frequencies. We note that our data suggest inappropriate functioning of this mechanism may contribute to the etiology of some diseases associated with p38 ^2,20–28^. As we find it is possible to elicit different genetic responses regardless of the precise nature of the stimulus, these findings may have implications for development of pharmaceutical and therapeutic strategies in a variety of diseases.

## Methods

### Data and Materials Availability

Microscopy data are available from the BioImage Archive (BIA, S-BIAD1827). RNAseq data are available from the Sequence Read Archive (SRA, BioProject PRJNA1225323) and Gene Expression Omnibus (GEO, GSE289999). PerKy-38 was provided by Dr. Taichiro Tomida. The plasmid pUC-GW-kan-VPC is available from Addgene (Addgene ID 244163) under MTA with UC Riverside. Plasmids pB510B-1 and pB210PA-1 for piggybac-mediated stable transfection are restricted from distribution under purchase agreement with Systems Biosciences. Code used for analysis and simulation is available at https://github.com/kuhlman-lab-ucr/p38.

### Plasmids, Constructs, and Proteins

PerKy-38 was the kind gift of Dr. Taichiro Tomida ^19^. The coding sequence for VPC, expressed from a CMV promoter, was designed using VectorNTI software (Thermo Fisher Scientific) and synthesized *de novo* and cloned into the high copy number pUC-GW-kan (GENEWIZ/Azenta). A second construct, VPC-2xStrep, was similarly designed and synthesized for VPC purification. VPC was then subcloned into the expression vector pB510B-1 (System Biosciences) by digestion of both pUC-GW-kan-VPC and pB510B-1 with NheI-HF and BamHI-HF (New England Biolabs), gel purification of the VPC band from pUC-GW-kan (Qiagen QIAquick gel extraction kit) and ligation into digested pB510B-1. The resulting plasmid, pB510B-1-VPC, was verified by nanopore sequencing (Plasmidsaurus).

SARS-CoV-2 Spike S1 subunit protein was purchased from R&D Systems (Minneapolis, MN). IL-1β was purchased from Invitrogen (Waltham, MA). TNF was purchased from InvivoGen (San Diego, CA). Sorbitol was purchased from Fisher Scientific. p38 inhibitor SB203580 was purchased from Selleck Chemicals (Houston, TX).

### Cell lines and culturing

Transient transfection of HEK-293T cells with pB510B-1-PerKy-38, pB510B-1-VPC, or pUC-GW-kan-VPC-2xStrep was performed using FuGENE HD Transfection Reagent (Promega) at a ratio of 3:1. HEK-293T cells stably transfected with PerKy-38 or VPC were generated by PiggyBac Transposon Vector System (System Biosciences) according to the manufacturer’s directions. Briefly, HEK-293T cells were doubly transfected with the pB510B-1 vectors along with PiggyBac Transposase vector pB210PA-1. Cells were allowed to recover for 48 hours, after which 5 µg/ml puromycin was added and cultures were expanded. After 14 days, surviving cells exhibiting blue and yellow fluorescence were collected, and the sequence of the inserted cassette verified by PCR and sequencing.

For experiments described here, cells were cultured in complete media containing 90% DMEM with high glucose (Gibco), 10% heat-inactivated FBS (Gibco), 200 mM L-glutamine (100X), 10 mM MEM nonessential amino acids (100X), 100 mM MEM sodium pyruvate (100X), and puromycin (Gibco). Phenol red pH indicator was omitted from the medium to avoid interfering with fluorescence imaging.

### Protein Purification and In Vitro FRET Assay

HEK-293T cells maintained as above were transiently transfected with pUC-GW-kan-VPC-2xStrep. Transfected cells were washed with ice-cold Dulbecco’s Phosphate Buffered Saline (DPBS), collected in a 15 ml tube (Falcon), and spun for 5 mins at 4 °C, 300*g*. The supernatant was discarded and cells were lysed by addition of 0.5 ml ice-cold Lysis buffer [1X RIPA buffer (Cell Signaling Technologies) + cOmplete Protease Inhibitor Cocktail (Roche) + Phosphatase Inhibitor Cocktail (EMD Millipore)]. Contents were transferred to a pre-chilled 1.5 ml Eppendorf tube and incubated on ice for 10 mins on a platform rotator. 3U per 10^6^ cells Benzonase nuclease (Sigma-Aldrich) was added, followed by another 15 min incubation on ice. The sample was centrifuged at 10,000*g*, 4 °C for 10 mins, and the supernatant collected into a 1.5 ml pre-chilled Eppendorf tube.

VPC-2xStrep protein was purified from this lysate using MagStrep Strep-Tactin XT beads (IBA Lifesciences) with a magnetic separator (IBA Lifesciences). After washing 2x with Wash buffer (100 mM Tris-HCl, 150 mM NaCl, 1 mM EDTA), protein was eluted 4x with magnetic beads elution buffer (100 mM Tris-HCl, 150 mM NaCl, 1 mM EDTA, 2.5 mM desthiobiotin). Pure protein was verified by sodium dodecyl sulfate polyacrylamide gel electrophoresis (SDS-PAGE) and western blotting (see below and **Extended Data Fig. 1**). VPC protein concentration was determined using the bicinchoninic acid (BCA) protein assay kit (Pierce/Thermo Fisher Scientific; Rockford, IL).

In vitro FRET assays were performed using VPC protein purified as above. VPC protein was diluted to a working concentration of 600 nM (20 mM HEPES, 100 mM NaCl). We performed FRET measurements on six experimental conditions: 1. Dilution buffer + 2.5 mM ZnCl_2_; 2. VPC (as purified) + 2.5 mM ZnCl_2_; 3. VPC + 0.25 mM ATP-Mg + 2.5 mM ZnCl_2_; 4. VPC + 0.25 mM ATP-Mg + 2.5 mM ZnCl_2_ + 100 U/ml Antarctic Phosphatase (New England Biolabs); 5. VPC + 0.25 mM ATP-Mg + 2.5 mM ZnCl_2_ + 200 nM MKK6 kinase (Sigma-Aldrich); and 6. VPC + 0.25 mM ATP-Mg + 2.5 mM ZnCl_2_ + 200 nM MKK6 kinase + 100 U/ml Antarctic Phosphatase. 100 μl of each reaction was prepared in the wells of a 96 well black optical plate (Thermo Fisher Scientific) and incubated at 37 °C for one hour. Samples were then measured in a BMG CLARIOstar Plus plate reader, with excitation at 430±10 nm and emission at 480±10 nm (mCerulean3) and 515±10 nm (mVenus). Each measurement was performed with three biological replicates with five technical replicates each. Values reported are the mean of these fifteen replicates, with error bars the standard error of the mean calculated as

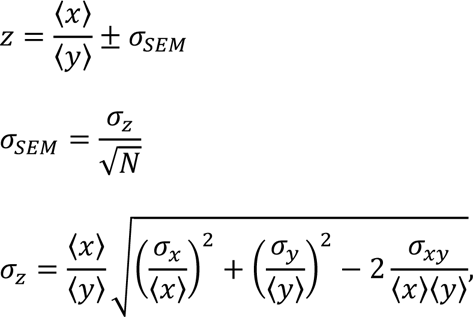

where 〈*x*_*i*_〉, σ_*xi*_, and σ_*xy*_ are the means, standard deviations, and covariances of measurements.

### Western Blot Analysis

For Western blot analysis, cells were grown as described above in 6 well plates (Corning Costar). Cells were washed with PBS and lysed with 100 μl ice cold Lysis buffer. This mixture was pulse sonicated to ensure complete lysis and then centrifuged at 21,000 *g* for 15 minutes in a refrigerated centrifuge and subsequently stored at -80 °C for use.

For Western blotting, 25µg of protein samples were mixed with 4x lithium dodecyl sulfate (LDS) sample buffer (Life Technologies) and 10x reducing agent (Invitrogen) and heated for 5 min at 100°C. These samples were loaded in a 4-12% SDS-PAGE gel (Mini-PROTEAN TGX, Bio-Rad) for electrophoretic separation and subsequently electro-transferred to nitrocellulose membrane (Bio-Rad). Membranes were blocked with 5 % bovine serum albumin (BSA; phospho blots; RPI Corp; 830075) or 5% milk (dry powdered milk; non-phospho blots; RPI Corp; M17200) in 1x TBST (tris buffered saline with Tween-20) for 1h at room temperature, washed 3x with 1x TBST for 5 min each, and afterward incubated overnight at 4°C in primary antibodies. We probed for fully activated p38 doubly phosphorylated at Thr180 + Tyr182 ^51^. The following primary antibodies (rabbit) were used: p38 (Cell Signaling Technologies; 9212S; 1:1000 1x TBST + 5% milk powder), phospho-p38 (ProSci; 9169; 1:1000 1x TBST + 5% BSA), and GAPDH (Cell Signaling Technologies; 2118S; 1:1000 1x TBST + 5% milk powder). Membranes were washed with 1x TBST (3 times for 15 min) and incubated with goat anti-rabbit-HRP (Invitrogen; 31460; 1:2000 1x TBST 1% BSA or milk powder as appropriate) secondary antibody. Blots were washed 3x with 1x TBST for 5 min each. Blots were stained by exposure to SuperSignal West Dura Extended Duration Substrate (Thermo Fisher).

Membranes were imaged with a Bio-Rad ChemiDoc system and images were quantified using Fiji/ImageJ ^52^. Values reported are the means of three biological replicates with error bars indicating the standard error of the mean. For those values reported as ratios, *e.g.*, p38/GAPDH, values and error bars were calculated as

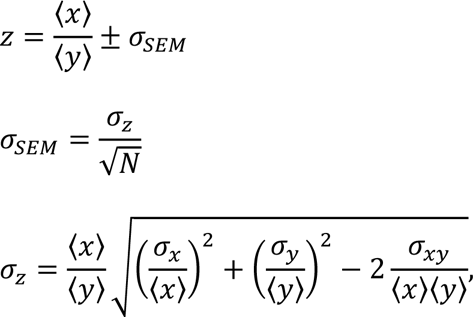

where 〈*x*_*i*_〉, σ_*xi*_, and σ_*xy*_ are the means, standard deviations, and covariances of measurements.

### Fluorescence Microscopy

Cell seeding for microscopy was done at 30% confluency in 35 mm glass cover slip dishes (Cellvis). Cells were incubated at 37°C and 5% CO_2_ in the glass cover slip dishes for at least 24 hours prior to imaging.

Cells were imaged using a motorized Nikon Ti-E Inverted Microscope and cultured in an Ibidi Silver Line stage top incubator at 37°C, 5% CO_2_, and 95% relative humidity. mCerulean3 was excited with a Nikon Intensilight epifluorescent illuminator using a D436/20 excitation filter (Chroma Technology), and both FRET reporters were imaged simultaneously using a Photometrics Dual View with YFP/CFP filter set on an Andor iXon DU897 Ultra camera. Images were taken using 500 ms exposure every 10 minutes scanned over a Field of View (FOV) grid programmed into the stage. For a single experiment, we typically imaged ∼100 FOVs over 48 hours, with an average of 7 – 8 cells per FOV.

Time traces of sinusoidally varying sorbitol concentration were quantified using media containing the fluorescent dye rhodamine B.

### Pump System and Dynamic Stimulation

Media flow was created using four New Era NE-9000 series peristaltic pumps with attached YG4 head kit facilitating 1/16^th^ inch inner diameter tubing. Food grade silicone tubing, 1/16^th^ inch inner diameter 1/8^th^ inch outer diameter, sanitized with 70% ethanol before use connects the pumps to the media bottles, mixing chamber, experiment dish, and waste bottle. The experiment dish was capped with a custom-made glass cover to allow insertion of tubing by a barbed elbow tubing joint. The pump rates were controlled individually using their RS-232 serial connectors from a custom-made python program using common serial, USB, and NumPy libraries to generate sinusoidal concentration waves of sorbitol over time.

### Image Segmentation and Cell Tracking

The single cell analysis workflow is summarized as: (1) pre-process images with FIJI/ImageJ ^52^ and segment cells with Cellpose ^53^; (2) stabilize identities with a prior-owner-aware Hungarian tracker that encodes motion, shape overlap, and size continuity; (3) align those identities with donor/acceptor frames; (4) estimate per-cell local background with a morphological ring; (5) compute background-corrected donor and acceptor signals with optional adaptive gating and bleed-through correction; and (6) derive per-cell FRET ratios and time series suitable for downstream statistical analysis. All computations were performed in Python using NumPy and pandas for data handling, tifffile for TIFF I/O, OpenCV for video rendering and text overlays, SciPy for optimization and morphology utilities, scikit-image for filtering and adaptive thresholding, and Matplotlib for visualization.

We first separated fluorescence channels of each image and performed background subtraction with FIJI/ImageJ. We then segmented each image to identify individual cells using Cellpose (v4.0.6) with GPU acceleration enabled and using the pretrained cyto3 model (*e.g.*, **Supplementary Movie 1**). Segmentation masks were generated and saved as TIFF files to serve as inputs to a custom single-cell tracking and quantification pipeline implemented in Python.

To obtain temporally stable cell identities, we developed a “sticky” tracker that combines recent-owner locking with assignment by the Hungarian algorithm. At each frame, putative objects were extracted from the Cellpose label image and represented by their centroid, area, and binary mask. Before global assignment, we performed a hard pre-assignment (“prior-owner lock”): if a detection substantially overlapped the most recent owner and was consistent with a constant-velocity prediction, the previous track ID was re-used, preventing ID swaps during transient overlaps, merges, or partial occlusions. The remaining objects and active tracks were matched by solving a linear assignment with a cost that blends four terms: predicted displacement relative to a constant-velocity extrapolation, one-minus-intersection over union (IoU) between current and dilated prior masks to favor shape continuity, an area-change penalty to discourage implausible growth/shrinkage, and a mild jump penalty for large inter-frame motions. We enforced a mutual-best IoU criterion to reduce asymmetric claims, applied a short memory window so tracks could persist through brief missed detections, and implemented a conservative revive step that re-attaches a new detection to a recently ended track if overlap and motion continuity are consistent. All masks were shape-checked and aligned defensively, and very small detections were ignored to suppress noise.

Single-cell FRET time series were obtained by pairing the stable track identities with their corresponding donor and acceptor image sequences. For each frame, the Cellpose-derived label of a given track ID was aligned with the donor and acceptor channels to ensure consistent spatial correspondence. A ring-shaped background region was generated by morphological dilation around each cell mask, and the median donor and acceptor intensities within this ring were used as local background references. The mean donor and acceptor intensities within each tracked cell were then measured, background-subtracted, and used to compute the FRET ratio as the quotient of corrected acceptor over donor signals. Frame indices were converted to elapsed time to create continuous single-cell trajectories. The pipeline exported per-track tables containing all quantitative features (frame, time, donor and acceptor signals, background-corrected values, and FRET ratios).

### Fourier Analysis

Details of implementation of Fourier analysis can be found in **SM I**. Before calculating the Fourier spectrum, we first subtracted the mean level of the FRET oscillations, setting the constant DC term of the Fourier spectrum to zero (see **SD V**). We then averaged the power spectra to generate the spectra presented in **Figs. 2,3**. The numbers of cells averaged was *N* ≥ 300 in all cases, with variation resulting from reliability of segmentation and tracking.

### RNAseq

Total RNA was extracted from HEK-293T cells using RNeasy Mini Kit (Qiagen) according to the manufacturer’s directions. All samples were collected as two independent biological replicates. This total RNA was provided to Novogene for strand specific messenger RNA (mRNA) sequencing. mRNA was purified from total RNA using poly-T oligo-attached magnetic beads. After fragmentation, first strand cDNA was synthesized using random hexamer primers, followed by second strand cDNA synthesis using dUTP for strand specific libraries. Libraries then underwent end repair, A-tailing, adapter ligation, size selection, amplification, and purification. Libraries were checked with Qubit and real-time PCR for quantification and bioanalyzer for size distribution detection.

After library quality control, different libraries were pooled based on the effective concentration and targeted data amount and sequenced by 150 bp paired end sequencing on an Illumina Novaseq X Plus platform. Raw reads in fastq format were first processed through fastp software. In this step, clean data (clean reads) were obtained by removing reads containing adapter, reads containing poly-N and low-quality reads from raw data. At the same time, Q20, Q30 and GC content of the clean data were calculated. All downstream analyses were based on high quality clean data, with each sample including more than 80 x 10^6^ reads. Reads were aligned to the hg38 *Homo sapiens* reference genome downloaded from NCBI. Index of the reference genome was built using Hisat2 v2.0.5 and paired-end clean reads were aligned to the reference genome using Hisat2 v2.0.5. featureCounts v1.5.0-p3 was used to count the read numbers mapped to each gene.

Differential expression (DE) analysis was performed using Matlab, which employs an implementation of the DESeq differential expression analysis pipeline ^54^. Each condition was divided by a library size factor equal to the median ratio of each feature over the geometric mean of the feature in all conditions to normalize the count prior to performing the hypothesis test. We identified genes resulting in a fold change relative to the negative control greater than 2x (*i.e.*, |log_2_(fold change)| ≥ 1) with adjusted *p*-value *p_adj_* ≤ 0.05 as being statistically significantly differentially expressed. Functional enrichment analysis was performed using g:Profiler (version e113_eg59_p19_f6a03c19) with g:SCS multiple testing correction method applying significance threshold of 0.1 ^55^. Data sources used for functional enrichment analysis include Gene Ontology ^56^ (GO) molecular function (GO:MF), GO cellular component (GO:CC), GO biological process (GO:BP), KEGG ^57^, Reactome (REACT) ^58^, WikiPathways (WP) ^59^, TRANSFAC (TF) ^60^, MiRTarBase (MIRNA) ^61^, Human Protein Atlas (HPA) ^62^, CORUM ^63^, and Human Phenotype Ontology (HP) ^64^.

## Acknowledgments

We thank Nigel Goldenfeld, Edward C. Cox, Hernan Garcia, Matthew Scott, Prue Talbot, Nathan Gabor, Joseph Rudnick, and Arthur Zhenyu Jia for useful discussions. This study was supported by startup funds provided by University of California, Riverside (TEK), UCR School of Medicine, Dean’s Collaborative Seed Grant (TEK, MK, RZ), University of California Office of the President UC Multicampus Research Programs and Initiatives grant M21PR3267 (TEK, MK, RZ), National Science Foundation, NSF DMR-2131963 (RZ), and National Institutes of Health NIH, R01 DA052209 (MK).

## Author Information

These authors contributed equally: Michael Worcester, Shayan Nejad.

## Author Contributions

SO, MK, RZ, and TEK conceived the study. TEK designed and oversaw all experiments and developed and wrote all theory. MW and SN performed microscopy and Fourier analysis. MW, SN, and TEK performed data visualization. DO designed, built, and operated the pump system. SA and AEN extracted RNA, prepared samples for RNAseq, and analyzed RNAseq data. PM purified proteins, performed westerns and SDS-PAGE, and PM and MG performed *in vitro* FRET assays. RZ and TEK oversaw simulations, and SL and KY performed simulations. MW, SN, AA, and NYY prepared cell lines and cell samples. TEK wrote the initial version of the paper, with review and revisions by all authors.

## Ethics Declarations

### Data and materials availability

Microscopy data are available from the BioImage Archive (BIA, S-BIAD1827). RNAseq data are available from the Sequence Read Archive (SRA, BioProject PRJNA1225323) and Gene Expression Omnibus (GEO, GSE289999). PerKy-38 was provided by Dr. Taichiro Tomida. The plasmid pUC-GW-kan-VPC is available from Addgene (Addgene ID 244163) under MTA with UC Riverside. Plasmids pB510B-1 and pB210PA-1 for piggybac-mediated stable transfection are restricted from distribution under purchase agreement with Systems Biosciences.

### Code availability

Code used for analysis and simulation is available at https://github.com/kuhlman-lab-ucr/p38.

## EXTENDED DATA FIGURES

**Extended Data Fig.1.**
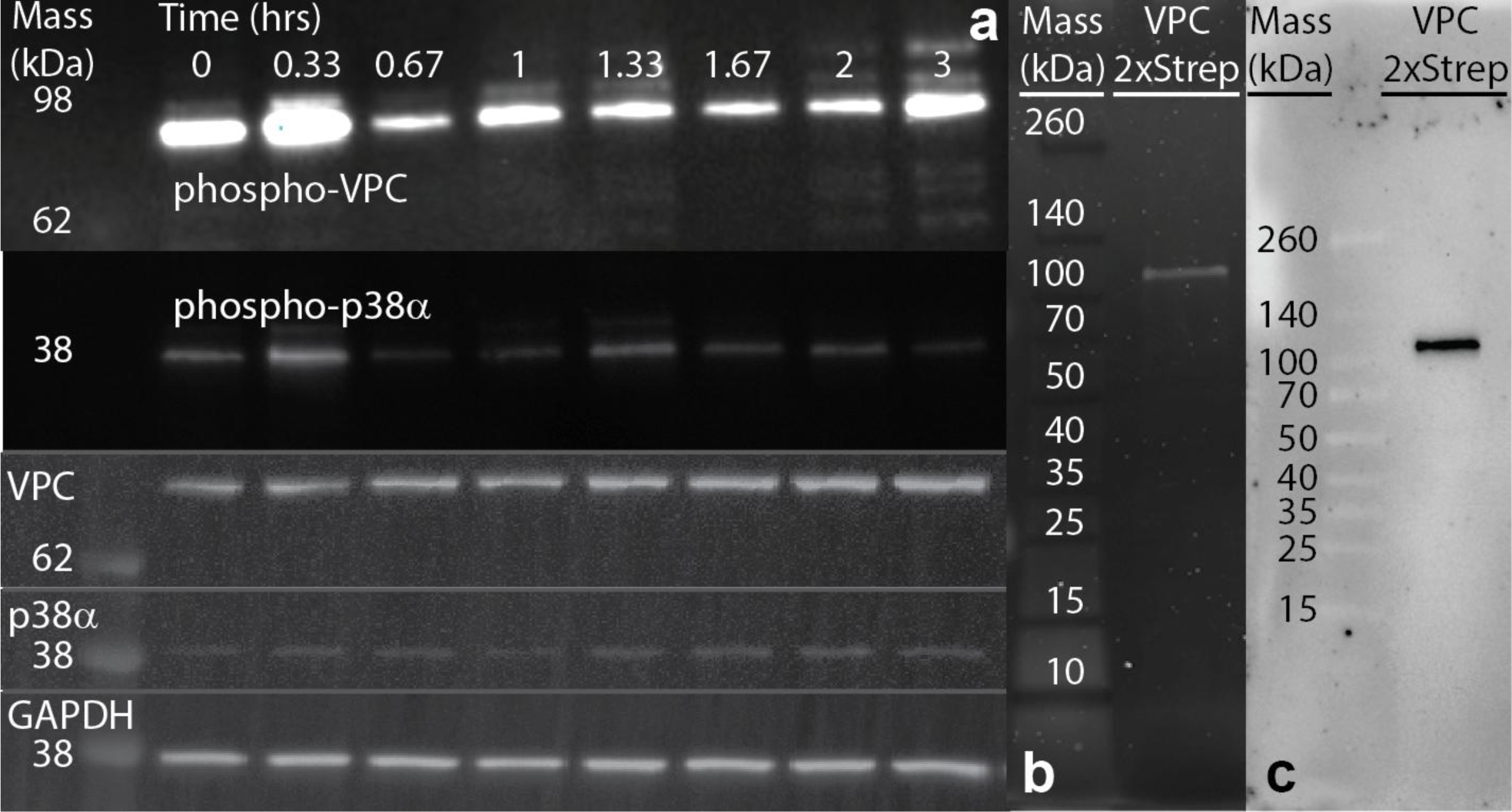
Western blot analysis of p38α and VPC phosphorylation. (a) Example western blots as a function of time after addition of 300 nM IL-1β of (from top): phospho-VPC, phospho-p38α; total VPC; total p38α; and GAPDH loading control. Each point in **Fig. 1c** is calculated as 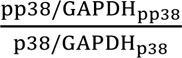 with three independent biological replicates; *i.e.*, each point includes data from 12 separate blots. (**b**) SDS-PAGE gel of purified VPC 2xStrep with SYPRO staining. (**c**) Western blot of purified VPC 2xStrep.

**Extended Data Fig. 2.**
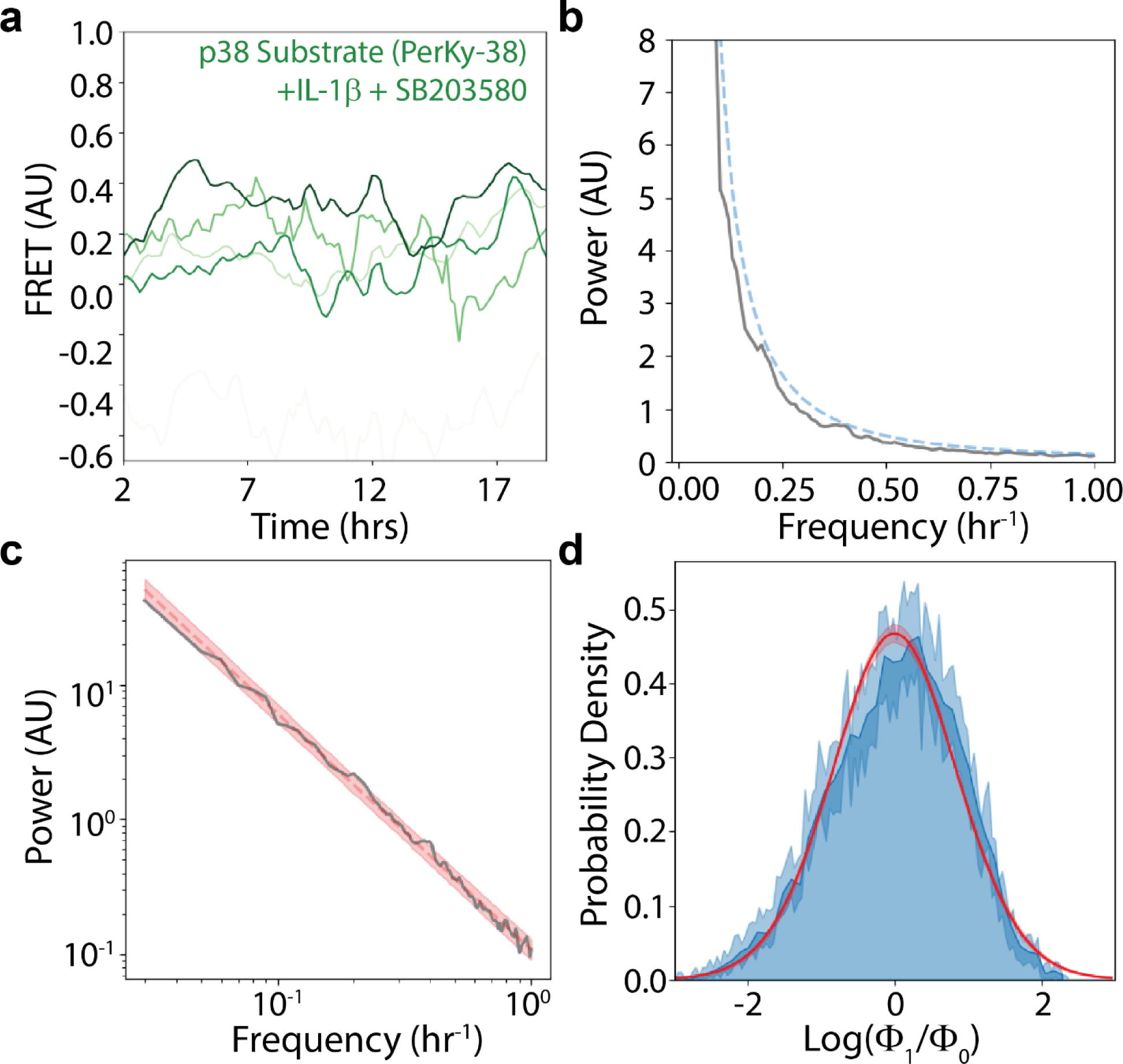
PerKy293+SB203580 Control Statistics. (a) Example FRET time traces of PerKy293 cells + 300 ng/mL IL-1β + 10 µM SB203580. (b) Average power spectrum of N = 450 PerKy293 cells as in (a). Blue dashed line: 99% confidence interval determined using statistics characterized below. (c) Log-log plot of PerKy293 + IL-1β + SB203580 power spectrum. Red dashed line: fit to *f* ^-*m*^ power law, yielding *m* = 1.74 ± 0.34, red shaded area shows best fit plus/minus error. (**d**) Histogram of deviations of PerKy293 + IL-1β + SB203580 power spectrum, Φ_1_, from power law fit, Φ_0_. Dark blue: deviations from best fit; shaded blue: deviations from best fit plus/minus errors shown in (b). Solid red line: best fit to log normal distribution. Shaded red region: error in fit to log normal distribution.

**Extended Data Fig. 3.**
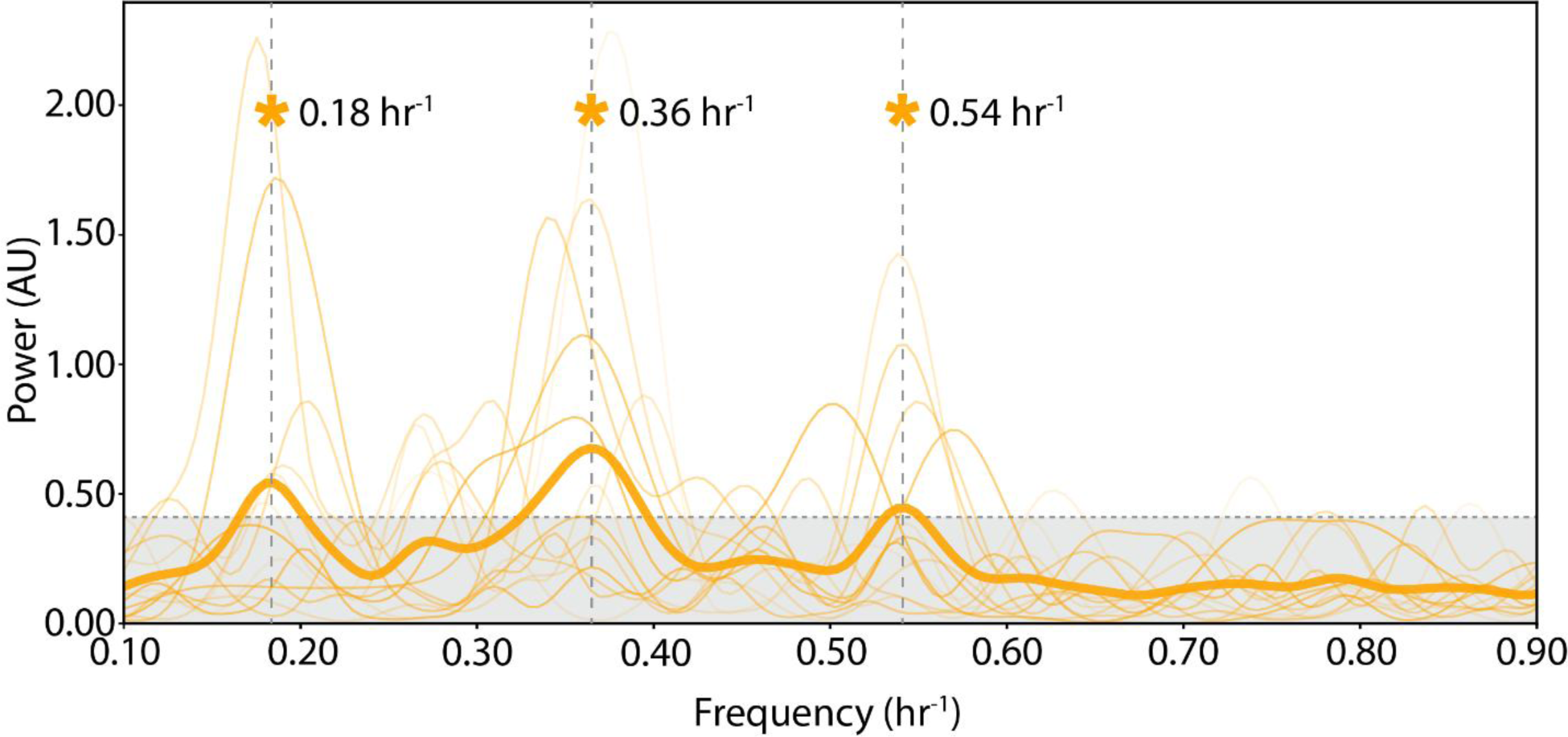
Averaging of Single Cell Spectra. Detrended (*i.e.*, noise power law spectrum subtracted) spectra of 14 example individual VPC293 cells stimulated with 300 ng/ml IL-1β (thin lines). Measurement and tracking noise (**Supplementary Movie 1**) introduce spurious peaks and errors in peak frequency locations that are suppressed in the average spectrum (thick line). Asterisks indicate statistically significant peaks in the average spectrum at the frequencies indicated by vertical dashed lines. Horizontal dashed line indicates 99% confidence bound as in **Fig. 2e-i**.

**Extended Data Fig. 4.**
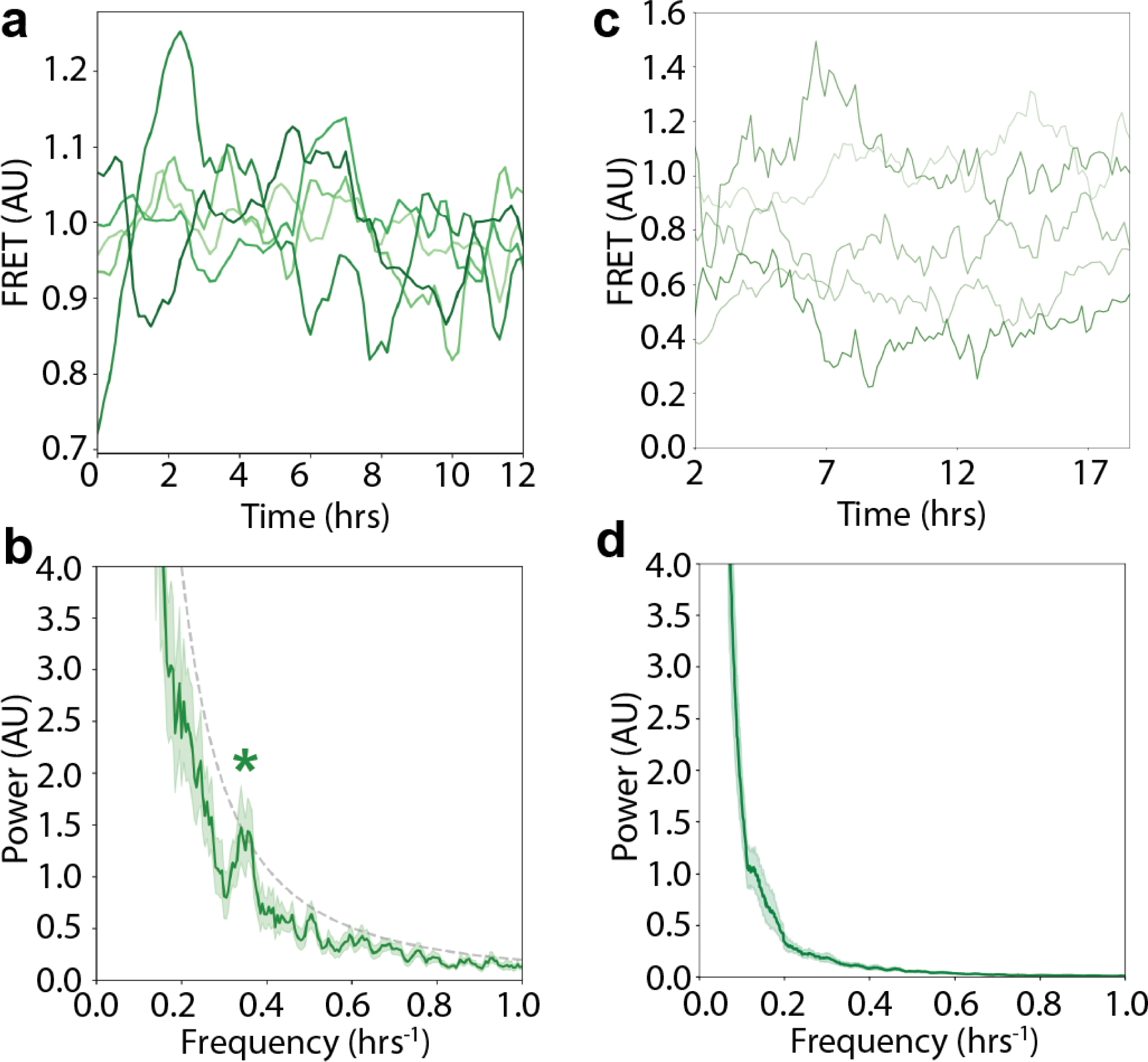
PerKy293 Response to Continuous Hyperosmotic Shock and TNF Stimulation. (**a**) FRET time traces of PerKy293 cells + 200 nM TNF. (**b**) Average power spectrum of *N* = 380 PerKy293 cells + 200 nM TNF. Note statistically significant peak (asterisk) at *f* = 0.36 hr^-1^, *p* = 2.9 x 10^-2^. (**c**) Example FRET time traces of PerKy293 cells + 120 mM sorbitol. (**d**) Averaged power spectrum of *N* = 600 PerKy293 cells + 120 mM sorbitol.

**Extended Data Fig. 5.**
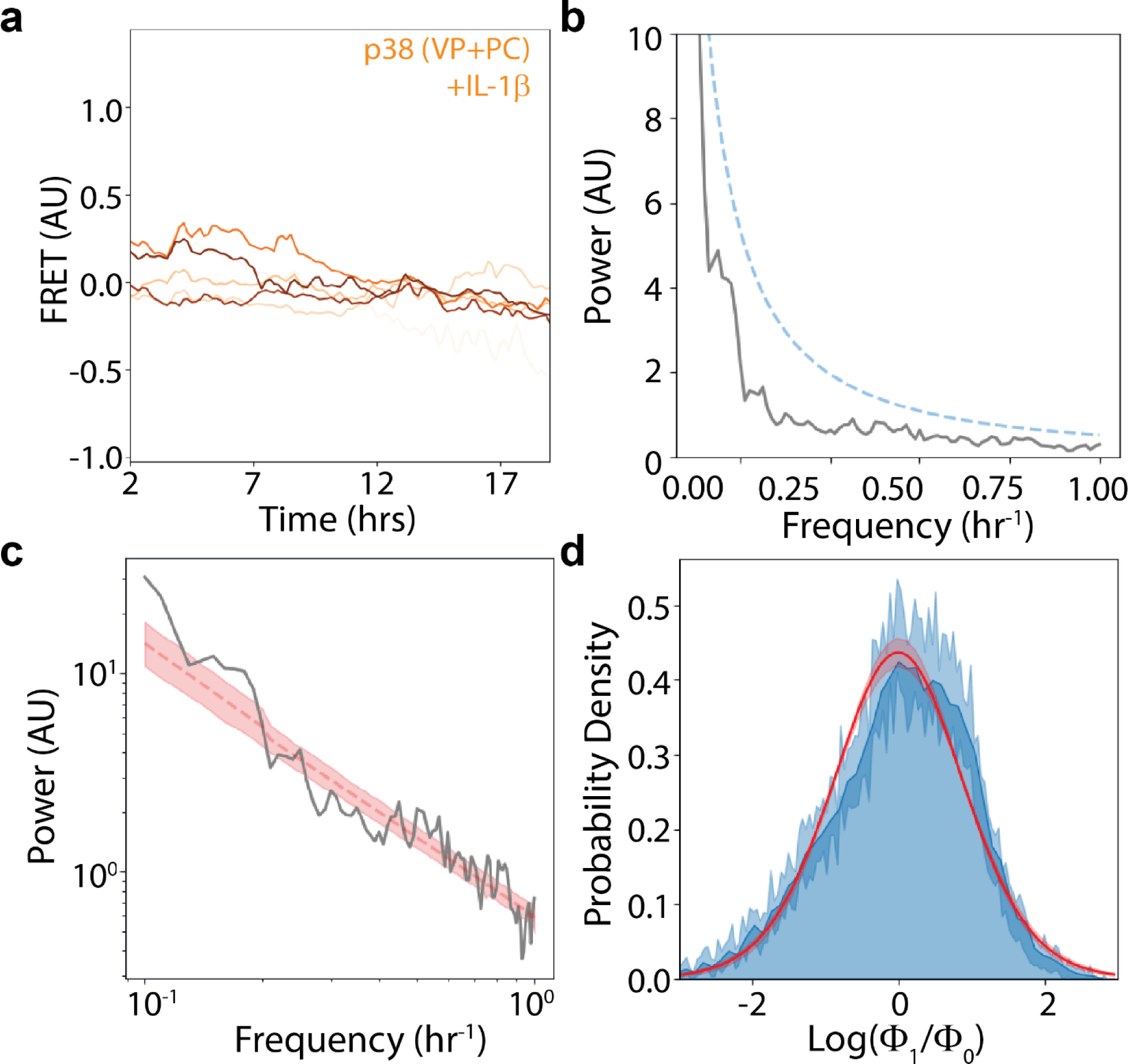
VPPC293 Control Statistics. (**a**) Example FRET time traces of VPPC293 control cells + 300 ng/mL IL-1β. (**b**) Average spectrum of *N* = 410 VPPC293 cells + IL-1β. Blue dashed line: 99% confidence interval determined using statistics characterized below. (**c**) Log-log plot of VPPC293 + IL-1β power spectrum. Red dashed line: fit to *f* ^-*m*^ power law, yielding *m* = 1.86 ± 0.20, red shaded area shows best fit plus/minus error. (**d**) Histogram of deviations of VPPC293 power spectrum, Φ_1_, from power law fit, Φ_0_. Dark blue: deviations from best fit; shaded blue: deviations from best fit plus/minus errors shown in (b). Solid red line: best fit to log normal distribution. Shaded red region: error in fit to log normal distribution.

**Extended Data Fig. 6.**
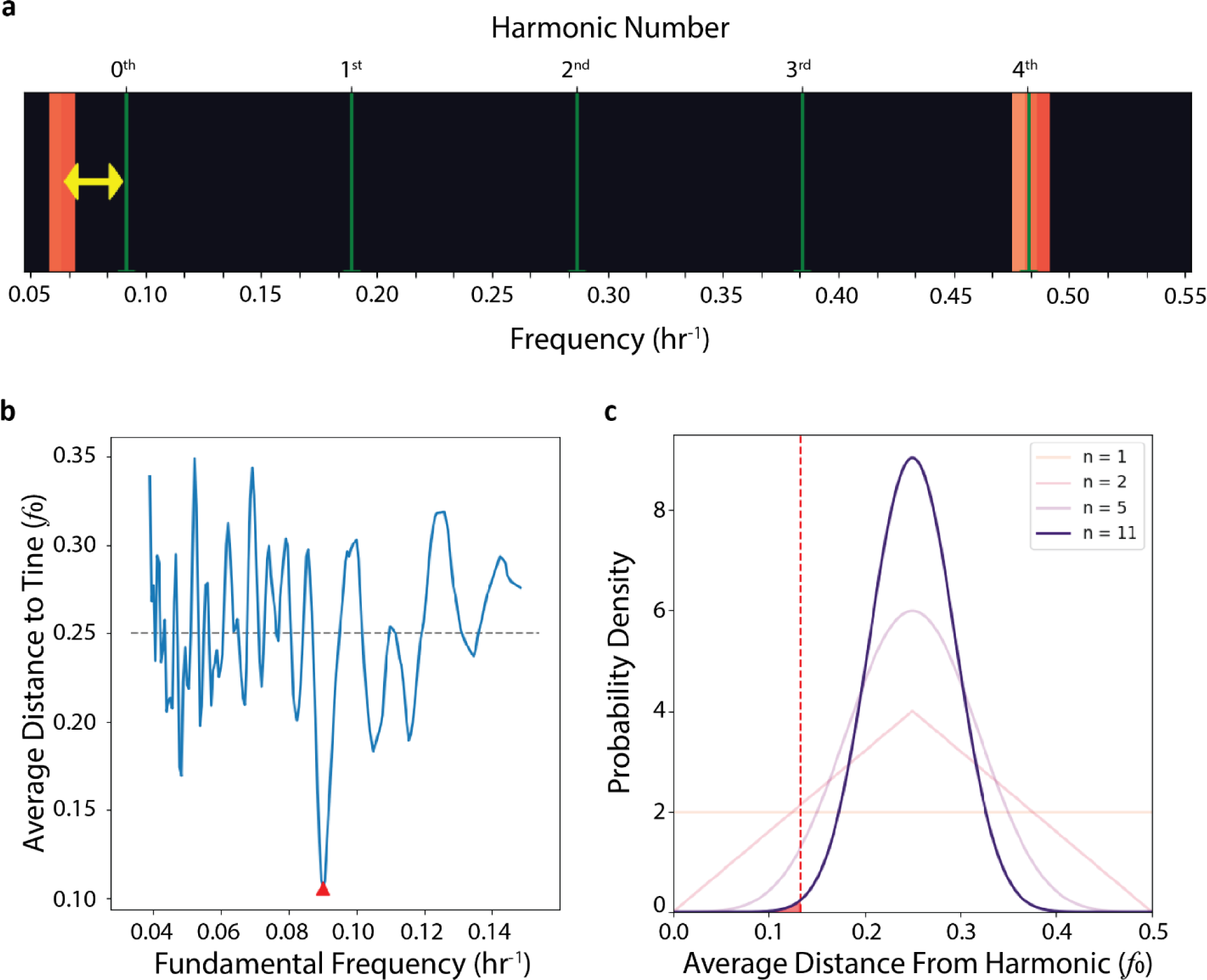
Harmonic Analysis of Stimulus Spectra. (**a**) The distances (yellow) of identified frequency bands (red, as in **Fig. 2k**) are measured relative to the tines of a Dirac comb with even harmonic spacing between the tines (green). (**b**) Average distance of frequency bands to the tines of the Dirac comb as a function of the fundamental frequency of the comb. The best fit fundamental value corresponds to minimizing this average distance (indicated by red), yielding *f*_0_ = 0.090 ± 0.002 hr^-1^. (**c**) Distribution of the mean distance of anharmonically (randomly) placed frequency bands from their nearest harmonic (*i.e.*, the Bates distribution on the interval [0,1/2]) in units of the fundamental frequency *f*_0_, demonstrated for different numbers of frequency bands. As we measure 11 frequency bands in the aggregate distribution (**Fig. 2K**, top), we use *n* = 11 (purple curve) to calculate *p* values, with distances measured from the center of mass of the band (red dashed line).

**Extended Data Fig. 7.**
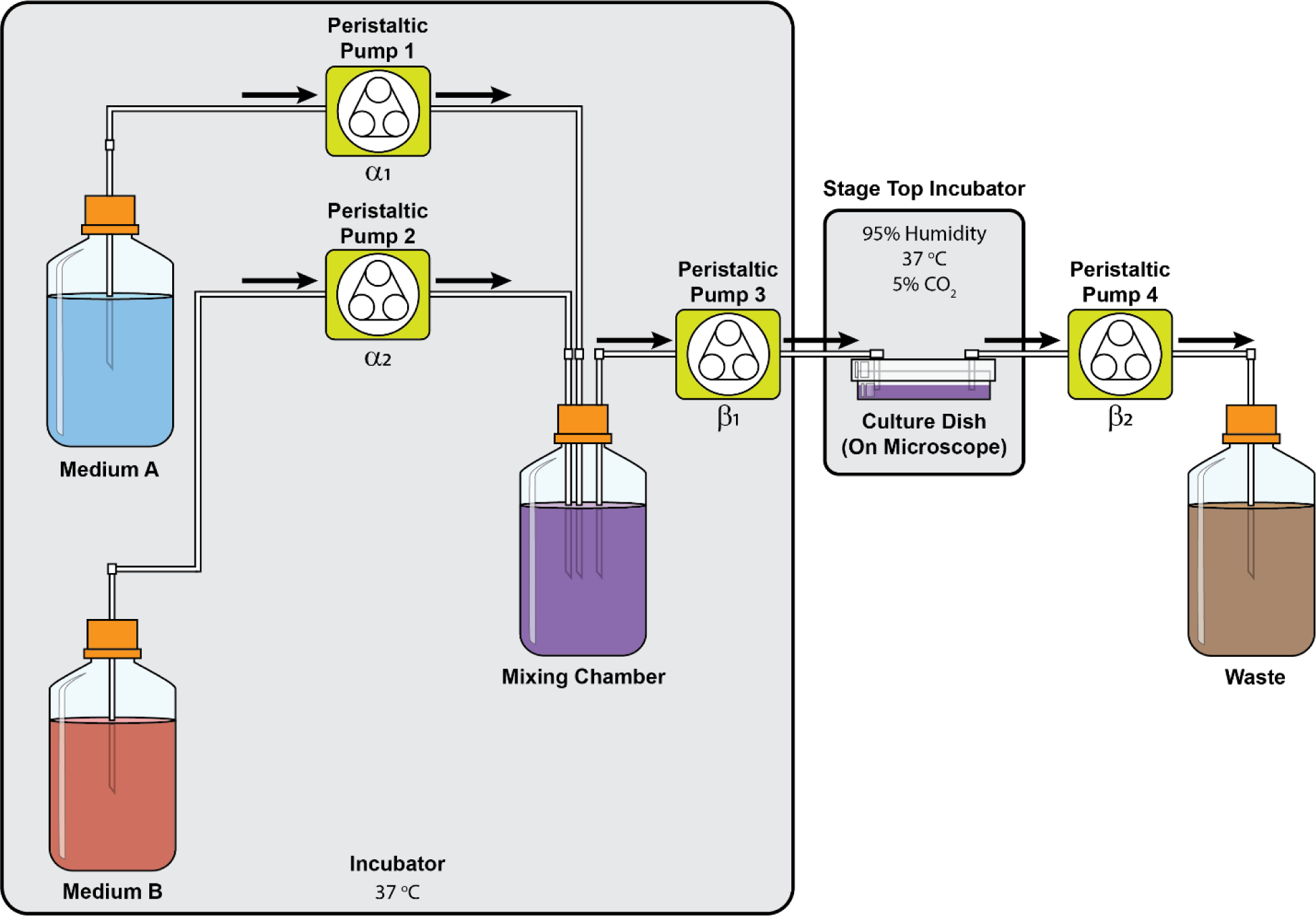
Pump System for Oscillatory Stimulation. Schematic illustration of the pump system used to deliver sinusoidally oscillating flows of sorbitol to cells within the microscope stage top incubator. The peristaltic pumps are connected by RS-232 connection to the PC controlling the microscope and running custom python software to control pump flow rates. The flow rates of each peristaltic pump (α_1_, α_2_, β_1_, and β_2_) are programmatically adjusted to deliver continuously varying sorbitol concentration at predetermined frequencies, with the constraints α_1_ + α_2_ = β_1_and β_1_ = β_2_ to maintain constant volumes.

**Extended Data Fig. 8.**
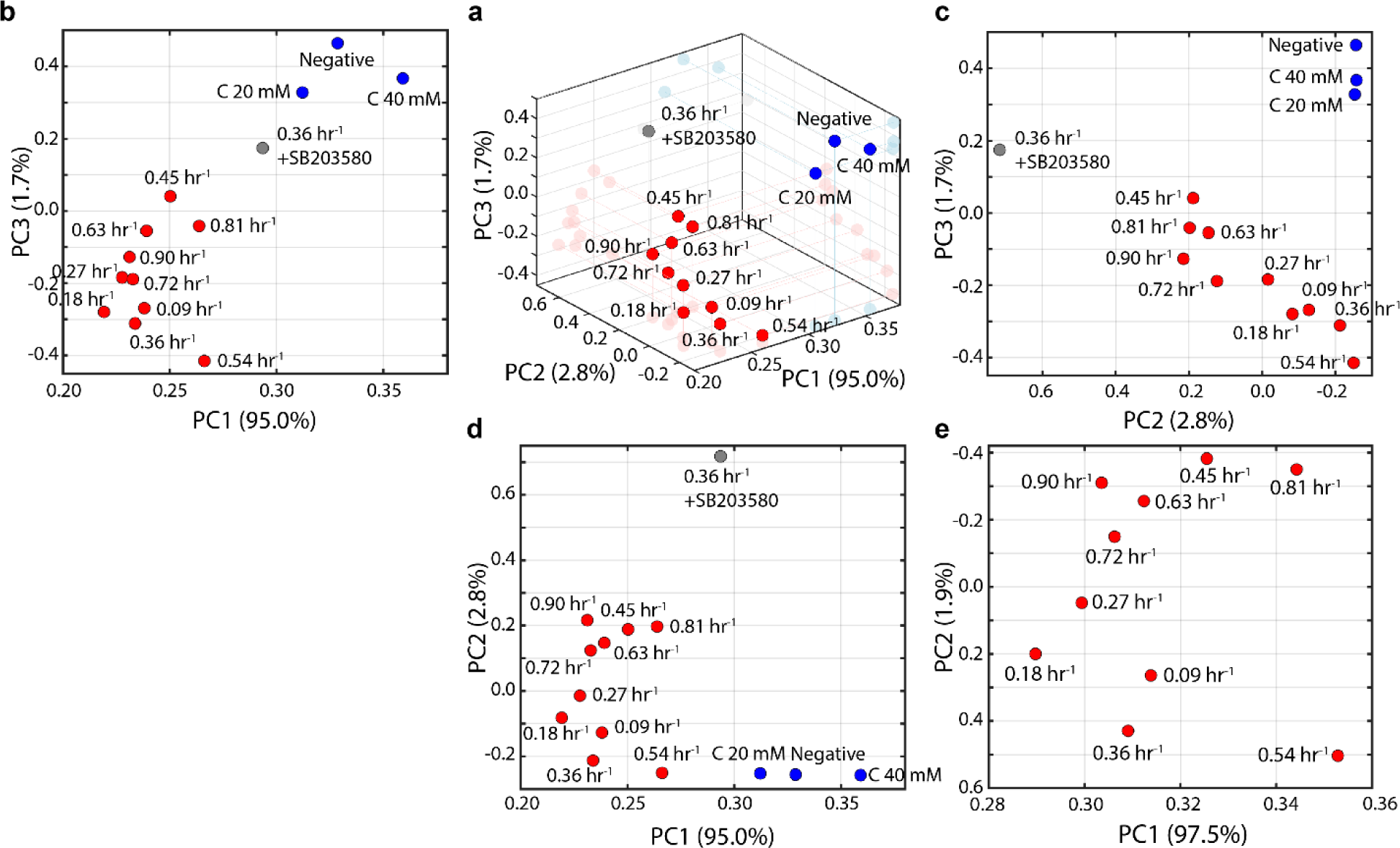
RNAseq Principal Component Analysis. (a) 3D plot of the principle components of RNAseq samples, identical to that shown in main text **Fig. 4**. (**b**-**d**) Projections of 3D data onto each 2D plane. (**e**) PC analysis of only periodically driven samples.

**Extended Data Fig. 9.**
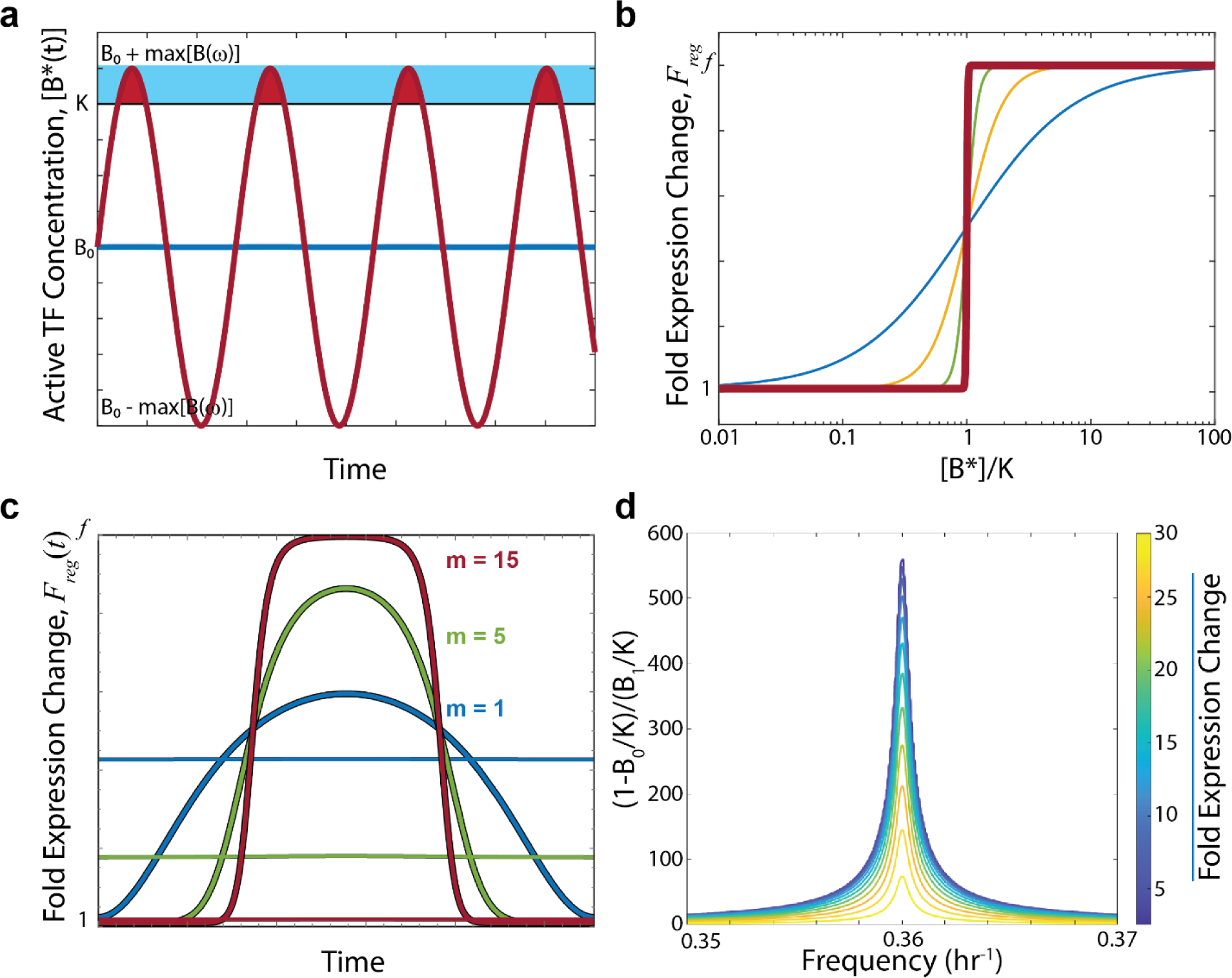
Bioresonant Transcriptional Regulation. (a) Concentration of active resonant (red) and off-resonant (blue) TF over time, eq. S7.4. The dissociation constant for binding to a *cis*-regulatory region, *K*, is indicated by a horizontal black line. The duty cycle, *D*(ω) (eq. S7.15), is the ratio of the red to blue shaded areas over one period. (b) The regulatory function *F_reg_* (eq. S7.3) plotted as a function of active TF concentration for various values of the Hill coefficient, *m*. Blue, *m* = 1; orange, *m* = 3; green, *m* = 10; red, *m* = 100. The “duty cycle” approximation corresponds to large values of *m* resulting in a switch-like step function, *e.g.*, the red curve. (**c**) The regulatory function *F_reg_* plotted as a function of time (*i.e.*, the integrand of eq. S7.7) for various values of the Hill coefficient, *m*, for resonant (thick lines) and off-resonant (thin lines) TFs. The ratio of areas under the thick versus thin lines represents the enhancement of gene expression when a TF is on-resonance versus off-resonance. Parameter values used: *f* = 100, *B*_0_/*K* = 0.73, *B*_1_/*K* = 0.001, *f*_0_ = 0.36 hr^-1^, γ = 0.004. (**d**) The gene expression fold change as a function of the driving frequency and parameter combination (1 − *B*_0_/*K*)/(*B*_1_/*K*) under the duty cycle approximation, eqs. S7.14 and S7.15. Note the narrow range of the x-axis, indicating that gene expression is enhanced only very close to the resonant frequency of the TF. Parameter values used: *f* = 100, *f*_0_ = 0.36 hr^-1^, γ = 0.004.

