## Supplementary Information for "Human Stress Response Specificity through Biochemical Resonance Selectivity"

**Supplementary Methods**

**SM I. Fourier Analysis**

The FRET signal from cells is a continuous function of time, $x(t)$. Our measurements are taken at discrete times with sampling period $T_s$ = 10 min, corresponding to a sampling frequency $f_s = \frac{1}{T_s}$. This yields a discrete sequence of $N$ FRET measurements $x_n = \{x_0, x_1, \ldots, x_{N-1}\}$. We obtain the frequency spectrum of the oscillations, $\tilde{X}_f$, as the discrete Fourier transform (DFT) of $x_n$,

$$\tilde{X}_f = \sum_{n=0}^{N-1} x_n e^{-i2\pi f n T}. \qquad \text{(S1.1)}$$

$\tilde{X}_f$ is a discrete spectrum of $N$ evenly spaced frequencies from $f = 0$ to $f = \frac{(N-1)}{NT} = \frac{(N-1)f_s}{N}$, *i.e.*,

$$\{f\} = \left\{0, \frac{f_s}{N}, \frac{2f_s}{N}, \ldots, \frac{(N-1)f_s}{N}\right\}. \qquad \text{(S1.2)}$$

The maximum frequency with unique information will not exceed the Nyquist frequency $f_{Nyquist} = f_s/2$, and hence we take $\tilde{X}_f$ as the subset of the DFT output containing $N/2$ elements with

$$\{f\} = \left\{0, \frac{f_s}{N}, \frac{2f_s}{N}, \ldots, \frac{f_s}{2}\right\}. \qquad \text{(S1.3)}$$

A complication is that, due to the discrete nature of $\tilde{X}_f$ and therefore its limited resolution, if the sampled frequencies do not correspond to the frequencies at which peaks occur in the continuous spectrum, these peaks will not be apparent in the discrete spectrum. To remedy this issue and increase the resolution with which the spectra are rasterized, we implement a procedure to analyze subsets of the available data points to generate a spectrum that contains samples of the discrete spectrum at more frequency points, $\tilde{g}_f$. We first generate a set of discrete spectra, $\tilde{X}_f^{(j)}$, with different numbers of data points, $j$, from the original data set. The number of data points in the set, $j$, runs from a minimum number of points $j_{min} = N/2 + 1$ to a maximum $j = N$,

$$\tilde{X}_f^{(j)} = \sum_{t=0}^{j} x_n e^{-i2\pi f t}, \qquad j = j_{min}, \ldots, N, \qquad \text{(S1.4)}$$

each of which yields output with a different set of frequencies.

We then collect these together as $\tilde{g}_f$,

$$\tilde{g}_f = \bigcup_{j=j_{min}}^{N} \tilde{X}_f^{(j)}. \qquad \text{(S1.5)}$$

### SM II. Statistical Significance of Fourier Peaks

We find that PerKy-38 and p38 activation state measured by FRET exhibit Brownian noise (*i.e.*, a power law $Af^m$ with $m$ = -2; **Extended Data Figs. 2, 5**), and therefore ensemble averaged Fourier spectra can be characterized as a power law function [**Extended Data Figs. 2c, 5c**]. The deviations of the data from the fitted line in log space are approximately normally distributed in our controls (**Extended Data Figs. 2d**, **5d**), enabling us to use normal statistics to generate $p$ values and confidence intervals.

To generate the line spectra shown in **Fig. 2k**, from each cell FRET time trace we first subtracted the mean FRET level and calculated the Fourier spectrum. We then removed the noise spectrum by subtraction in log space. We identified the frequencies where all local maxima occur, defined as any points of magnitude higher than their nearest neighbors. The only constraint applied is that adjacent peaks could not be closer than 0.02 $hr^{-1}$, a value determined through visual inspection of the behavior of multiple individual spectra. Each local maximum was subsequently weighted by multiplying by the power of the respective frequency bin in the Fourier spectrum, with the rationale that high power peaks should contribute more strongly than those with low power. We then built a histogram of the weighted counts of peaks at each frequency with bin width 0.005 $hr^{-1}$. The histogram built in such a way from unstimulated VPC293 cells is shown in **Fig. 2j**, with the resulting normal distribution of the magnitude of deviations of counts from the mean obtained by integrating over all frequencies shown to the right of **Fig. 2j**. The spectra of those frequencies where peaks are over-represented in cells subjected to various stimuli are shown in **Fig. 2k**, with statistical significance (*i.e.*, $z$ score) calculated using the distribution derived as described above and shown in **Fig. 2J**, with a cutoff threshold set at $z = 2$.

### SM III. Analysis of Harmonics

Stimulus line spectra as in **Fig. 2k** display statistically over-represented frequencies in Fourier spectra. We analyzed these line spectra to describe the likelihood of the spectra being harmonically organized with evenly spaced frequencies, and to determine the fundamental frequency from which the harmonics are derived.

For a given fundamental frequency $f_0$, we generate a series of harmonics as a Dirac comb, $Ш_{f_0}(f)$,

$$Ш_{f_0}(f) = \sum_{n=1}^{\infty} \delta(f - nf_0)\,, \tag{S3.1}$$

represented by the green tines in **Extended Data Fig. 6a**. We then measure the distance of each frequency band in the aggregate spectrum shown in **Fig. 2k** to the nearest harmonic; the distance of a frequency band can be located no more than $f_0/2$ from the closest tine of the Dirac comb. The best fit fundamental frequency is determined by scanning all possible $f_0$ and minimizing the average distance-to-tine (**Extended Data Fig. 6b**). Using this method, we find the best fit fundamental frequency to be $f_0 = 0.090 \pm 0.002$ $hr^{-1}$.

However, a series of bands placed at random frequencies may artifactually appear harmonic when judged by their distance to the nearest harmonic. Assuming the frequencies where bands appear are random, the probability of finding a single band a given distance from the nearest tine of the Dirac comb is a uniform distribution over the interval $[0, f_0/2]$. Consequently, the probability to find $n$ frequency bands an average distance $X$ from their closest tine is the Bates distribution on the interval [0, 1/2], *i.e.*, the distribution of the mean, $X$, of $n$ independent random variables distributed uniformly over [0, 1/2]:

$$f_X(x;n) = \frac{n}{(n-1)!}\sum_{k=0}^{n}(-1)^k\binom{n}{k}(2nx-k)^{n-1}\text{sgn}(2nx-k)\,, \tag{S3.2}$$

where $\binom{n}{k}$ is the binomial coefficient

$$\binom{n}{k} = \frac{n!}{k!\,(n-k)!} \tag{S3.3}$$

and

$$\text{sgn}(2nx-k) = \begin{cases} -1 & 2nx < k \\ 0 & 2nx = k\,. \\ 1 & 2nx > k \end{cases} \tag{S3.4}$$

As we observe 11 frequency bands in our aggregated spectrum, we calculate $p$ values using this distribution with $n = 11$ and $x$ measured from the center of mass of the band, as shown in **Extended Data Fig. 6c**.

**Supplementary Discussion and Equations**

**SD IV. Forced Oscillations, Resonance, and Resonance Selectivity**

A linear harmonic oscillator with viscous damping driven by a sinusoidally oscillating force, $F(t)$ [*e.g.*, $F(t) = F_0\cos(\omega t)$], is described by

$$\ddot{x} + \gamma\dot{x} + \omega_0^2 x = F(t), \tag{S4.1}$$

where $x(t)$ is the time-dependent displacement of the oscillator, $\gamma$ the damping parameter, and $\omega_0$ is the natural angular frequency ($\omega_0 = 2\pi f_0$) of the undamped oscillator. $F_0$ and $\omega$ are the coupling strength and angular frequency of the driving oscillator, respectively. Dot notation indicates time derivatives.

Taking the Fourier transform,

$$-\omega^2\tilde{x}(\omega) - i\omega\gamma\tilde{x}(\omega) + \omega_0^2\tilde{x}(\omega) = \tilde{F}(\omega), \tag{S4.2}$$

and solving for $\tilde{x}(\omega)$

$$\tilde{x}(\omega) = \frac{1}{\omega_0^2 - \omega^2 - i\gamma\omega}\tilde{F}(\omega) \tag{S4.3.1}$$

$$= \frac{\omega_0^2 - \omega^2 + i\gamma\omega}{(\omega_0^2 - \omega^2)^2 + (\gamma\omega)^2}\tilde{F}(\omega). \tag{S4.3.2}$$

This defines the transfer function, $\chi(\omega)$ (see **SD VI**),

$$\tilde{x}(\omega) = \chi(\omega)\tilde{F}(\omega), \tag{S4.4.1}$$

$$\chi(\omega) = \frac{\omega_0^2 - \omega^2}{(\omega_0^2 - \omega^2)^2 + (\gamma\omega)^2} + i\frac{\gamma\omega}{(\omega_0^2 - \omega^2)^2 + (\gamma\omega)^2}. \tag{S4.4.2}$$

The long-time solution after decay of transients is found by taking the inverse Fourier transform

$$x(t) = |\chi(\omega)|e^{i\Phi(\omega)}F(t). \tag{S4.5.1}$$

The frequency dependent amplitude response of the driven oscillator is given by the modulus of the transfer function

$$|\chi(\omega)| = \frac{1}{[(\omega_0^2 - \omega^2)^2 + (\gamma\omega)^2]^{1/2}} \tag{S4.5.2}$$

see **SD VI** below and **Fig. 3**. The phase difference between driver and target is given by

$$\Phi(\omega) = \tan^{-1}\left[\frac{\gamma\omega}{\omega_0^2 - \omega^2}\right]. \tag{S4.6}$$

The resonant frequency, $\omega_r$, is found from the oscillatory part of the poles of the transfer function,

$$\omega_r = \sqrt{\omega_0^2 - \left(\frac{\gamma}{2}\right)^2}\,. \tag{S4.7}$$

If damping is weak ($\gamma \ll 1$), $\omega_r \cong \omega_0$. Consequently, the amplitude of the driven oscillations peaks when $\omega \cong \omega_0$, corresponding to resonance between driver and target.

Resonance selectivity occurs when a driving oscillator with angular frequency $\omega$ is coupled to multiple targets, each with their own natural angular frequencies $\omega_{0,j}$. When the driving frequency is close to the natural frequency of one of the targets, $\omega \sim \omega_{0,a}$, the amplitude of the oscillations of target $a$ is selectively amplified relative to those targets with natural frequencies not near that of the driver $\omega_{0,j}$ ($j \neq a$).

The Quality factor, $Q$, is defined relative to the full width at half maximum (FWHM) of the amplification peak, $\Delta\omega$,

$$Q \equiv \frac{\omega_0}{\Delta\omega} = \frac{\omega_0}{\gamma}. \tag{S4.8}$$

The relative magnitude of the amplification at resonance is characterized by $Q$,

$$\frac{|\chi(\omega = \omega_0)|}{|\chi(\omega = 0)|} = Q. \tag{S4.9}$$

Hence, ideal resonance selectivity with large, selective amplification of specific targets corresponds to a system with large $Q$ factor.

**SD V. Biochemical Resonance**

We demonstrate that any two-component biological oscillator, when coupled to an oscillating driving enzyme, will demonstrate all the familiar phenomena of driven damped oscillators, including resonance and resonance selectivity, as described above.

The binding of an enzyme, $E$, to a substrate, $S$, to produce a product, $P$, is described by the reaction

$$E + S \underset{k_{-1}}{\overset{k_1}{\leftrightarrow}} ES \underset{k_{cat}}{\longrightarrow} E + P, \tag{S5.1}$$

with $k_{cat}$ the catalytic rate constant and $k_1$and $k_{-1}$ the forward and reverse binding rate constants, respectively. The rate of product formation is

$$\frac{dP}{dt} = k_{cat}\frac{ES}{K + S}\,, \tag{S5.2}$$

with $E$ the enzyme concentration, $S$ the substrate concentration, and $K$ the Michaelis constant.

If the substrate is in excess, then $S >> K$ and the rate increases linearly with the enzyme concentration

$$\frac{dP}{dt} \cong k_{cat}E. \tag{S5.3}$$

Alternatively, if the amount of substrate is low, then $S << K$ and the rate of product formation depends on both $E$ and $S$,

$$\frac{dP}{dt} \cong \frac{k_{cat}}{K}ES. \tag{S5.4}$$

While resonance between driver and target will occur in all regimes, the effect of bioresonance is simplest and most clearly seen in the linear case where all species are present in excess, *i.e.*, the regime described by eq. S5.3. A typical model for a two-component biological oscillator, *e.g.*, the p53/Mdm2 system [35], in the linear regime is given as

$$\frac{dx_1}{dt} = -\beta_1 x_1 - \alpha_2 x_2 \tag{S5.5.1}$$

$$\frac{dx_2}{dt} = \alpha_1 x_1 - \beta_2 x_2, \tag{S5.5.2}$$

with $x_j$ the concentration of each component, $\alpha_j$ the catalytic rates, and $\beta_j$ the rates of spontaneous dephosphorylation/degradation.

In matrix notation,

$$\begin{bmatrix} \dot{x}_1 \\ \dot{x}_2 \end{bmatrix} = \begin{bmatrix} -\beta_1 & -\alpha_2 \\ \alpha_1 & -\beta_2 \end{bmatrix} \begin{bmatrix} x_1 \\ x_2 \end{bmatrix} \tag{S5.6.1}$$

$$\dot{\mathbf{x}}(t) = \mathbf{A}\mathbf{x}(t). \tag{S5.6.2}$$

If this system is driven enzymatically by a third component, $z(t)$, with catalytic rates $\zeta_1$ and $\zeta_2$, and if the unphosphorylated substrates, $x_j'$, are in excess relative to $z$ (*e.g.*, eq. S5.3), then

$$\begin{bmatrix} \dot{x}_1 \\ \dot{x}_2 \end{bmatrix} = \begin{bmatrix} -\beta_1 & -\alpha_2 \\ \alpha_1 & -\beta_2 \end{bmatrix} \begin{bmatrix} x_1 \\ x_2 \end{bmatrix} + \begin{bmatrix} \zeta_1 z(t) \\ \zeta_2 z(t) \end{bmatrix} \tag{S5.7.1}$$

$$\dot{\mathbf{x}}(t) = \mathbf{A}\mathbf{x}(t) + \mathbf{z}(t), \tag{S5.7.2}$$

where $\zeta_j$ is the catalytic rate of the enzyme $z$ reacting with unphosphorylated substrate $x_j'$.

Taking the Fourier transform and solving for $\tilde{\mathbf{x}}(\omega)$,

$$(-i\omega\mathbf{I} - \mathbf{A})\tilde{\mathbf{x}}(\omega) = \tilde{\mathbf{z}}(\omega), \tag{S5.8}$$

$$\tilde{\mathbf{x}}(\omega) = (-i\omega\mathbf{I} - \mathbf{A})^{-1}\tilde{\mathbf{z}}(\omega) \tag{S5.9.1}$$

$$= \frac{\mathrm{adj}[-i\omega\mathbf{I} - \mathbf{A}]}{\det[-i\omega\mathbf{I} - \mathbf{A}]}\tilde{\mathbf{z}}(\omega) \equiv \mathbf{Y}(\omega)\tilde{\mathbf{z}}(\omega). \tag{S5.9.2}$$

This defines the Y-parameter admittance matrix $\mathbf{Y}(\omega)$ (see **SD VI**, eq. S6.5 - S6.10), given by

$$\mathbf{Y}(\omega) = \frac{\mathrm{adj}[-i\omega\mathbf{I} - \mathbf{A}]}{\det[-i\omega\mathbf{I} - \mathbf{A}]} = \frac{1}{\det[-i\omega\mathbf{I} - \mathbf{A}]}\begin{bmatrix} \beta_2 - i\omega & -\alpha_2 \\ \alpha_1 & \beta_1 - i\omega \end{bmatrix}. \tag{S5.10.1}$$

With the identifications

$$\omega_0^2 = \alpha_1\alpha_2 + \beta_1\beta_2 \tag{S5.10.2}$$
$$\gamma = \beta_1 + \beta_2, \tag{S5.10.3}$$

$\mathbf{Y}(\omega)$ is expressed as

$$\mathbf{Y}(\omega) = \frac{[(\omega_0^2 - \omega^2) + i\gamma\omega]}{[(\omega_0^2 - \omega^2)^2 + (\gamma\omega)^2]}\begin{bmatrix} \beta_2 - i\omega & -\alpha_2 \\ \alpha_1 & \beta_1 - i\omega \end{bmatrix} \tag{S5.10.4}$$

$$= \chi(\omega)\begin{bmatrix} \beta_2 - i\omega & -\alpha_2 \\ \alpha_1 & \beta_1 - i\omega \end{bmatrix}. \tag{S5.10.5}$$

Continuing from eq. S5.9.2,

$$\tilde{\mathbf{x}}(\omega) = \frac{1}{\det[-i\omega\mathbf{I} - \mathbf{A}]}\begin{bmatrix} \beta_2 - i\omega & -\alpha_2 \\ \alpha_1 & \beta_1 - i\omega \end{bmatrix}\begin{bmatrix} \zeta_1\tilde{z}(\omega) \\ \zeta_2\tilde{z}(\omega) \end{bmatrix}, \tag{S5.11.1}$$

or

$$\det[-i\omega\mathbf{I} - \mathbf{A}]\tilde{\mathbf{x}}(\omega) = \begin{bmatrix} \beta_2 - i\omega & -\alpha_2 \\ \alpha_1 & \beta_1 - i\omega \end{bmatrix}\begin{bmatrix} \zeta_1\tilde{z}(\omega) \\ \zeta_2\tilde{z}(\omega) \end{bmatrix}. \tag{S5.11.2}$$

Taking the inverse Fourier transform,

$$\ddot{x}_1 + \gamma\dot{x}_1 + \omega_0^2 x_1 = (\beta_2\zeta_1 - \alpha_2\zeta_2)z + \zeta_1\dot{z} \tag{S5.12.1}$$
$$\ddot{x}_2 + \gamma\dot{x}_2 + \omega_0^2 x_2 = (\beta_1\zeta_2 + \alpha_1\zeta_1)z + \zeta_2\dot{z}. \tag{S5.12.2}$$

Defining the driving chemical potentials, $\mu_j(t)$,

$$\mu_j(t) = \sum_k Y_{jk} z_k\,, \tag{S5.12.3}$$

these become the standard equations of forced, damped linear harmonic oscillators, eq. S4.1,

$$\ddot{x}_1 + \gamma\dot{x}_1 + \omega_0^2 x_1 = \mu_1(t) \tag{S5.12.4}$$
$$\ddot{x}_2 + \gamma\dot{x}_2 + \omega_0^2 x_2 = \mu_2(t). \tag{S5.12.5}$$

The resonant frequency of the system is found from the characteristic equation

$$-\omega_r^2 - i\gamma\omega_r + \omega_0^2 = 0, \tag{S5.13.1}$$

resulting in

$$\omega_r = \sqrt{\omega_0^2 - \left(\frac{\gamma}{2}\right)^2} \tag{S5.13.2}$$

$$= \sqrt{\alpha_1\alpha_2 - \left(\frac{\beta_1 - \beta_2}{2}\right)^2}\,. \tag{S5.13.3}$$

If damping is weak, $\beta_1, \beta_2, \gamma \sim 0$, and

$$\omega_r \cong \sqrt{\alpha_1\alpha_2}. \tag{S5.13.4}$$

To be consistent with the typical identification of $\omega_0$ as the undamped natural frequency of the system, we can alternatively define

$$\omega_0 = \sqrt{\alpha_1\alpha_2} \tag{S5.14.1}$$
$$\Gamma\ = 2i\sqrt{\beta_1\beta_2}. \tag{S5.14.2}$$

Then the natural angular frequency of the damped system, eq. S5.10.2, which we now call $\omega_0'$, is

$$\omega_0'^2 = \omega_0^2 - \left(\frac{\Gamma}{2}\right)^2\,, \tag{S5.14.3}$$

and the resonant frequency is expressed as

$$\omega_r = \sqrt{\omega_0^2 - \frac{\gamma^2 + \Gamma^2}{4}} \cong \omega_0 \tag{S5.14.4}$$

With these definitions, the appearance of $\Gamma$ in eq. S5.12 is similar to structural damping, where the damping force is proportional to displacement, $x(t)$, and typically considered in phase with the velocity, $\dot{x}(t)$, leading to hysteretic energy loss. However, because here $\Gamma^2$ is in phase with the displacement, the only effect is shifting the natural frequency. Throughout we will use $\omega_0^2 = \alpha_1\alpha_2 + \beta_1\beta_2$ for notational simplicity.

Now consider the required form of solutions to eqs. S5.12. To ensure that protein concentrations are always positive, the input $z$ and output solutions $x_j$ must be expressed as a Fourier spectrum with some positive constant offset [the "direct current (DC)" term of the Fourier spectrum],

$$x_j(t) = \overline{x_j} + \int d\omega\, \widetilde{x_j}(\omega)e^{i\omega t} \tag{S5.15.1}$$

$$z(t) = \bar{z} + \int d\omega\, \tilde{z}(\omega)e^{i\omega t} \tag{S5.15.2}$$

Substituting these into eqs. S5.12 and taking the time average, any oscillatory terms will average to zero, yielding

$$\omega_0^2\overline{x_1} = (\beta_2\zeta_1 - \alpha_2\zeta_2)\bar{z} \tag{S5.16.1}$$

$$\omega_0^2\overline{x_2} = (\beta_1\zeta_2 + \alpha_1\zeta_1)\bar{z}, \tag{S5.16.2}$$

or

$$\overline{x_1} = \frac{(\beta_2\zeta_1 - \alpha_2\zeta_2)}{\omega_0^2}\bar{z} \tag{S5.16.3}$$

$$\overline{x_2} = \frac{(\beta_1\zeta_2 + \alpha_1\zeta_1)}{\omega_0^2}\bar{z}. \tag{S5.16.4}$$

That is, when considering the driving potentials

$$\mu_1(t) = (\beta_2\zeta_1 - \alpha_2\zeta_2)z + \zeta_1\dot{z} \tag{S5.17.1}$$

$$\mu_2(t) = (\beta_1\zeta_2 + \alpha_1\zeta_1)z + \zeta_2\dot{z}, \tag{S5.17.2}$$

the first term, proportional to $z$, provides a direct current (DC) input that determines the average activation level of the substrate, $\overline{x_j}$. The second term, proportional to $\dot{z}$, corresponds to an alternating current (AC) input that drives oscillations about the average activation level. Intracellular protein levels here are measured through fluorescence, which introduces an unknown proportionality factor between fluorescence level and protein numbers. We therefore cannot extract any meaningful fitting parameters from absolute fluorescence or FRET values without further calibration. Hence, when calculating Fourier spectra, we first subtracted the mean level of the FRET oscillations, effectively setting the first term to zero and yielding

$$\mu_j = \zeta_j\dot{z}. \tag{S5.18}$$

As described in **SD IV**, ideal resonance selectivity requires a large $Q$ factor,

$$Q = \frac{\omega_0}{\gamma} = \frac{\sqrt{\alpha_1\alpha_2 + \beta_1\beta_2}}{\beta_1 + \beta_2} \; ; \tag{S5.19}$$

that is, a weakly damped system with small spontaneous degradation rates, $\beta_j$, relative to the catalytic rates, $\alpha_j$. However, for protein levels to always be positive, it must be the case that

$$\min\left[x_j(t)\right] = \bar{x}_j - \frac{\zeta_j}{\gamma}\min[z(t)] \geq 0. \tag{S5.20.1}$$

This sets a limit on the maximum possible $Q$ factor

$$Q = \frac{\omega_0}{\gamma} \leq \frac{\left(\beta_k\zeta_j \mp \alpha_k\zeta_k\right)}{\omega_0\zeta_j}\frac{\bar{z}}{\min[z(t)]}. \tag{S5.20.2}$$

While here we consider p38 MAPK and its substrates, where oscillations occur in the number of phosphorylated and unphosphorylated active/inactive molecules, this mathematical description of bioresonance is agnostic to the mechanistic nature of the oscillations. Bioresonance can be employed when oscillations of any nature are present, regardless of whether the oscillations occur in phosphorylation levels or some other mechanism, such as direct expression and degradation causing oscillations in total protein numbers.

### SD VI. Linear Response Theory and the p38/PerKy-38 Transfer Function

Here we demonstrate that the transfer function quantifies all aspects of the interaction between target and driver oscillators, including phase shifts, transfer of energy, and the amplitude response of the target. Hence characterization of a system's transfer function offers a concise and complete way to test hypotheses about the relationship between two oscillators.

The spectral response of a forced oscillator, $\tilde{x}(\omega)$, can be expressed as proportional to the spectrum of the driving input, $\tilde{\mu}(\omega)$, using the transfer function, $\chi(\omega)$, also referred to as the generalized susceptibility, as

$$\tilde{x}(\omega) = \chi(\omega)\tilde{\mu}(\omega). \tag{S6.1}$$

For a driven damped linear simple harmonic oscillator, the transfer function is given by eq. S4.4,

$$\chi(\omega) = \frac{(\omega_0^2 - \omega^2)}{[(\omega_0^2 - \omega^2)^2 + (\gamma\omega)^2]} + i\frac{\gamma\omega}{[(\omega_0^2 - \omega^2)^2 + (\gamma\omega)^2]}. \tag{S6.2.1}$$

$$\equiv \chi'(\omega) + i\chi''(\omega). \tag{S6.2.2}$$

For weak damping, $\gamma \ll 1$, only behavior near resonance is significant, $\omega \approx \omega_0$, yielding the Lorentzian form

$$\chi(\omega) \cong \frac{1}{2\omega_0}\left[\frac{(\omega_0-\omega)}{[(\omega_0-\omega)^2+(\gamma/2)^2]}+i\frac{\gamma/2}{[(\omega_0-\omega)^2+(\gamma/2)^2]}\right] \tag{S6.2.3}$$

The amplitude of driven oscillations is given by the modulus of the transfer function, $|\chi(\omega)|$ (see eqs. S4.5). The imaginary part of the transfer function, $\chi''(\omega)$, describes the time-averaged rate of energy absorption by the driven oscillator as work, $W$,

$$\overline{\frac{dW}{dt}} = \frac{\omega}{2\pi}\int_0^{2\pi/\omega} dt\left[\iint_{-\infty}^{\infty}\frac{d\omega}{2\pi}\frac{d\omega'}{2\pi}[-i\omega\chi(\omega)]e^{-i(\omega+\omega')t}F(\omega)F(\omega')\right] = 2F_0^2\omega\chi''(\omega), \tag{S6.3.1}$$

where $F_0$ is the coupling parameter between the driver and target (see. **SD IV** eq. S4.1). The phase shift between driver and target can be expressed using the transfer function as

$$\Phi(\omega) = \tan^{-1}\left(\frac{\chi''(\omega)}{\chi'(\omega)}\right) = \tan^{-1}\left[\frac{\gamma\omega}{\omega_0^2-\omega^2}\right]. \tag{S6.3.2}$$

If the spectra of both input and output are measured, the transfer function can be calculated easily according to eq. S6.1 as

$$\chi(\omega) = \frac{\tilde{x}(\omega)}{\tilde{\mu}(\omega)}. \tag{S6.4}$$

Comparison of the expected transfer function of a driven linear simple harmonic oscillator to that obtained experimentally for p38/PerKy-38 is complicated by the biological oscillator model, eqs. 5.7, employing matrices as variables rather than a single input and output. In electrical network theory, such a model is described as a two-port network with input voltages $z_j(t)$ generating output currents $x_k(t)$, where the Y-parameter admittance matrix, $\mathbf{Y}(\omega)$ plays a similar role as the transfer function; compare eq. S6.1 to eq. S5.9.2.

From eq. S5.9.2

$$\tilde{\mathbf{x}}(\omega) = \mathbf{Y}(\omega)\tilde{\mathbf{z}}(\omega) \tag{S6.5.1}$$

or

$$\tilde{x}_j(\omega) = \sum_k Y_{jk}(\omega)\tilde{z}_k(\omega). \tag{S6.5.2}$$

The expected PerKy-38 spectrum, $\tilde{x}_1(\omega)$ [or $\tilde{x}_2(\omega)$, the choice is arbitrary], is therefore given by

$$\tilde{x}_1(\omega) = \sum_k Y_{1k}(\omega)\tilde{z}_k(\omega) \tag{S6.6.1}$$

$$= \chi(\omega)[(\beta_2 - i\omega)\zeta_1\tilde{z}(\omega) - \alpha_1\zeta_2\tilde{z}(\omega)]. \tag{S6.6.2}$$

Collecting terms,

$$\tilde{x}_1(\omega) = -\omega\zeta_1[-\chi''(\omega) + i\chi'(\omega)]\tilde{z}(\omega) + (\beta_2\zeta_1 - \alpha_1\zeta_2)[\chi'(\omega) + i\chi''(\omega)]\tilde{z}(\omega). \quad \text{(S6.7)}$$

As described above (see **SD V**, eqs. S5.15 – S5.18), subtraction of the mean FRET values before calculating the Fourier spectrum effectively sets the second term to zero, yielding

$$Y_1(\omega) = -i\omega\zeta_1\chi(\omega) \quad \text{(S6.8.1)}$$

$$= -\omega\zeta_1[-\chi''(\omega) + i\chi'(\omega)]. \quad \text{(S6.8.2)}$$

This results in switching the real and imaginary parts and inverting the real part, as experimentally observed (**Fig. 3a,b)**. Therefore, the AC behavior of $x_i(t)$ behaves as

$$x_1(t) = -i\omega\chi(\omega)\zeta_1 z(t). \quad \text{(S6.9)}$$

If driven by a sinusoidal oscillator $z(t)$ with frequency $\omega$ (*i.e.*, p38), the amplitude of the driven oscillations in $x_1(t)$ (*i.e.*, PerKy-38) is given by the modulus of $Y_1(\omega)$,

$$|x_1(t)| = |Y_1(\omega)||z(t)| \quad \text{(S6.10.1)}$$

$$= \frac{\omega\zeta_1}{\sqrt{(\omega_0^2 - \omega^2)^2 + (\gamma\omega)^2}}|z(t)|, \quad \text{(S6.10.2)}$$

$$\cong \frac{\zeta_1/2}{\sqrt{(\omega_0 - \omega)^2 + (\gamma/2)^2}}|z(t)|, \quad \text{(S6.10.3)}$$

*cf.*, eq. S4.5 above. The modulus of $Y_1(\omega)$ for the p38/PerKy-38 system is shown in **Fig. 3c**, yielding a best fit value of $\gamma = 0.00423 \pm 0.00009$ (SEM) $hr^{-1}$. This corresponds to a quality factor, $Q \equiv \omega_0/\gamma$, of $Q = 535 \pm 11$. Again, since measurement by fluorescence introduces an unknown scaling due to arbitrary fluorescence units, $\zeta_1$ cannot be determined from these data by fitting.

### SD VII. Bioresonant Transcriptional Regulation

We next address whether bioresonance can provide specificity in gene regulation. We demonstrate that bioresonance can be used to specifically control genetic responses only if regulation of those genes is cooperative. This is consistent with experimentally observed generality of high cooperativity in eukaryotic transcriptional regulation [65].

A simple model of the output of a genetic circuit regulated by the binding and unbinding of protein transcription factors (TFs) at some *cis*-regulatory region can be described using statistical mechanics [66]. As the oscillations we observe have periods of multiple hours, we assume that on the short time scales of TF binding and unbinding the system can be treated as in equilibrium.

Consider a TF $B$ that can exist in either an active ($B^*$) or inactive ($B$) state. When active, it binds to a *cis*-regulatory region with dissociation constant $K$ and Hill coefficient $m$, which quantifies the degree of TF binding cooperativity. When bound, the TF interacts with RNA polymerase to a degree quantified by the capacity *f*. If RNAP polymerase binds at the promoter with dissociation constant $K_P$, the probability to find RNA polymerase bound at the promoter is described by [66]

$$P_{RNAP\ bound} = \frac{1}{1 + \frac{K_P}{[RNAP]F_{reg}}}, \tag{S7.1}$$

where $F_{reg}$ is given by

$$F_{reg} = \frac{1 + f \cdot \left(\frac{[B^*]}{K}\right)^m}{1 + \left(\frac{[B^*]}{K}\right)^m}. \tag{S7.2}$$

The rate of transcription, and therefore rate of gene expression, is proportional to this probability [66]. With the standard approximation that the promoter is weak, $[RNAP]/K_P \ll 1$,

$$\begin{aligned} P_{RNAP\ bound} &\cong \frac{[RNAP]}{K_P} F_{reg} \\ &= \frac{[RNAP]}{K_P} \cdot \frac{1 + f \cdot \left(\frac{[B^*]}{K}\right)^m}{1 + \left(\frac{[B^*]}{K}\right)^m}. \end{aligned} \tag{S7.3}$$

TFs that are activated and deactivated by phosphorylation and dephosphorylation reactions can result in oscillations of the TF activation state (see **SD V**). The amount of active TF at any time, *t*, is given by

$$B^*(\omega, t) = B_0 + B(\omega)\cos(\omega t + \varphi), \tag{S7.4}$$

with $B_0$ the mean level of activation, $\omega$ the angular frequency, and $\varphi$ an arbitrary phase shift. The amplitude of oscillations, $B(\omega)$, is given by eq. S4.5,

$$B(\omega) = \frac{\omega B_1}{[\gamma^2\omega^2 + (\omega_0^2 - \omega^2)^2]^{1/2}}, \tag{S7.5}$$

with $B_1$ a scaling parameter. For such a system, $\max[B(\omega)] = B_1/\gamma \leq B_0$ so that TF concentration always remains positive. Furthermore, for bioresonance to work as a transcriptional regulatory strategy, it must be the case that (**Extended Data Fig. 9a**)

$$B_0, \max[B(\omega)] \; < \; K \; \sim \; B_0 + \max[B(\omega)]. \tag{S7.6}$$

The time-averaged rate of gene expression is obtained as

$$\overline{\text{Gene Expression Rate}} \propto \overline{P_{RNAP\ bound}(\omega)} \cong \frac{1}{T}\frac{[RNAP]}{K_P}\int_0^T \frac{1 + f \cdot \left(\frac{[B^*(\omega,t)]}{K}\right)^m}{1 + \left(\frac{[B^*(\omega,t)]}{K}\right)^m} dt. \quad \text{(S7.7)}$$

For an off-resonant TF, $B(\omega) \cong 0$ and the integrand of eq. S7.7 loses time dependence. The integral eq. S7.7 can be performed trivially to yield

$$\overline{\text{Gene Expression Fold Change}_{OFF}} \equiv \overline{GEFC_{OFF}} = \frac{1 + f \cdot \left(\frac{[B_0]}{K}\right)^m}{1 + \left(\frac{[B_0]}{K}\right)^m} \equiv \frac{1 + f \cdot {b_0}^m}{1 + {b_0}^m}, \quad \text{(S7.8)}$$

*i.e.*, off-resonance TFs controlled by the same master regulator are differentially expressed to a degree quantified by $b_0 = [B_0]/K$.

For an on-resonance TF, eq. S7.7 can be evaluated analytically using, *e.g.*, Cauchy's residue theorem. However, the result quickly becomes unwieldy for $m > 1$. We demonstrate below that there must be some minimum degree of cooperativity characterized by a Hill coefficient $m^* > 1$ for bioresonance to specifically differentially regulate genetic responses.

For $m = 1$, we obtain from integrating eq. S7.7

$$\overline{GEFC(\omega)} = \frac{1 + fb_0\sqrt{1 - \left[\frac{b(\omega)}{1 + b_0}\right]^2}}{(1 + b_0)\sqrt{1 - \left[\frac{b(\omega)}{1 + b_0}\right]^2}} - f \cdot \frac{\left(1 - \sqrt{1 - \left[\frac{b(\omega)}{1 + b_0}\right]^2}\right)}{(1 + b_0)\sqrt{1 - \left[\frac{b(\omega)}{1 + b_0}\right]^2}} \quad \text{(S7.9)}$$

$$= \frac{1}{\delta(\omega)}\left[\frac{1 + \big(f \cdot \delta(\omega)\big)b_0}{(1 + b_0)} - f \cdot \frac{\big(1 - \delta(\omega)\big)}{(1 + b_0)}\right], \quad \text{(S7.10)}$$

with

$$\delta(\omega) \equiv \sqrt{1 - \left[\frac{b(\omega)}{(1 + b_0)}\right]^2} \leq 1 \quad \text{(S7.11)}$$

where we have again absorbed $K$ into the definitions of $b_0$ and $b_1$. Note that eqs. S7.9 – S7.10 reduce to eq. S7.8 when $b(\omega \neq \omega_0) \cong 0$ and $\delta(\omega \neq \omega_0) \cong 1$.

On resonance, $\omega = \omega_0$, and

$$\overline{GEFC_{ON}} = \frac{1}{\delta}\left[\frac{1+(f\cdot\delta)b_0}{(1+b_0)} - f\cdot\frac{(1-\delta)}{(1+b_0)}\right], \quad \delta = \sqrt{1-\left[\frac{2b_1}{\gamma(1+b_0)}\right]^2} \tag{S7.12}$$

For any reasonable choice of parameters (eq. S7.6), it is therefore the case when $m = 1$ that

$$\overline{GEFC_{ON}} \leq \overline{GEFC_{OFF}}, \tag{S7.13}$$

*i.e.*, bioresonance cannot provide specificity in transcriptional regulation when TF binding is non-cooperative ($m = 1$).

Conversely, for $m \gg 1$, we consider a duty cycle approximation of eq. S7.7. If TF binding at the *cis*-regulatory region is cooperative (*i.e.*, $m \gg 1$), $F_{reg}$ can be approximated as a step function switch; that is, expression is at its maximum level if $[B^*] > K$, whereas expression is completely off if $[B^*] < K$ (**Extended Data Fig. 9b**). With this approximation, the time-averaged gene expression fold change is

$$\overline{GEFC} \cong f\cdot D(\omega), \tag{S7.14}$$

where $D(\omega)$ is the TF's duty cycle, or the fraction of time when $[B^*] > K$ and the TF is bound. For sinusoidal oscillations of $B^*$ as given by eq. S7.4, $D(\omega)$ can be defined piecewise as

$$D(\omega) = \begin{cases} 0 & \dfrac{1-b_0}{b(\omega)} > 1 \\[2ex] \dfrac{2\sqrt{1-\left(\dfrac{1-b_0}{b(\omega)}\right)^2} - \left(\dfrac{1-b_0}{b(\omega)}\right)\left[\pi - 2\sin^{-1}\left(\dfrac{1-b_0}{b(\omega)}\right)\right]}{2\pi\left(\dfrac{1-b_0}{b(\omega)}\right)} & \dfrac{1-b_0}{b(\omega)} \leq 1 \end{cases}. \tag{S7.15}$$

Consequently, since $b(\omega) \cong 0$ when off-resonance, $D(\omega) \cong 0$, resulting in low expression. Conversely, $b(\omega) \gg 0$ for on-resonance TFs, resulting in an extremely sharp enhancement of gene expression only when $\omega = \omega_0$, as shown in **Extended Data Fig. 9d**.

We therefore conclude that for on-resonance TFs to lead to a greater effect on gene expression than off-resonant TFs, nonlinearity in TF binding must be present to suppress the level of gene expression when $[B^*] < K$ and amplify the effect when $[B^*] > K$ (**Extended Data Fig. 9b**). That is, TF binding at *cis*-regulatory regions must be cooperative for bioresonance to provide specificity in gene regulation (**Extended Data Fig. 9c-d**).

### SD VIII. Simulations of p38 bioresonance selectivity

For simulations of p38 activity shown in **Fig. 5**, we use the set of nonlinear differential equations describing p38 (MAPK) phosphorylation state developed by Tomida *et al.*[19],

$$\frac{d[\text{MAP2K}]}{dt} = k_0 S[t](1-[\text{MAP2K}]) - k_1[\text{MAP2K}] \tag{S8.1.1}$$

$$\frac{d[\mathrm{MAPK}]}{dt} = k_2[\mathrm{MAP2K}](1-[\mathrm{MAPK}]) - k_3[\mathrm{MKP1}]\frac{[\mathrm{MAPK}]}{k_4+[\mathrm{MAPK}]} \quad \text{(S8.1.2)}$$

$$\frac{d[\mathrm{MKP1RNA}]}{dt} = k_5[\mathrm{MAPK}](1-[\mathrm{MKP1RNA}]) - k_6[\mathrm{MKP1RNA}] \quad \text{(S8.1.3)}$$

$$\frac{d[\mathrm{MKP1}]}{dt} = k_7[\mathrm{MKP1RNA}](1-[\mathrm{MKP1}]) - k_8[\mathrm{MKP1}]. \quad \text{(S8.1.4)}$$

Parameter values are as given in Tomida *et al.*[19], multiplied by a factor of 0.1175 to yield the experimentally observed p38 fundamental frequency of 0.09 $\mathrm{hr}^{-1}$:

$$k_2 = 1.06\ \mathrm{hr}^{-1}$$
$$k_3 = 1.13\ \mathrm{hr}^{-1}$$
$$k_4 = 0.000705$$
$$k_5 = 0.38775\ \mathrm{hr}^{-1}$$
$$k_6 = 0.353\ \mathrm{hr}^{-1}$$
$$k_7 = 1.41\ \mathrm{hr}^{-1}$$
$$k_8 = 0.141\ \mathrm{hr}^{-1}$$

Note that the first equation for the time evolution of [MAP2K] is decoupled from and can be solved independently of the others. Tomida *et al.* used $S[t] = 1$ when p38 is activated by MAP2K and $S[t] = 0$ otherwise. Instead, we treat [MAP2K] as a driving oscillator representing p38's interactions with oscillating components of the phosphorylation cascade in the equation for the time evolution of [MAPK] (p38), setting $[\mathrm{MAP2K}] = A[1 + Bsin(2\pi ft)]$ with $A = 0.2857$ and $B$ a random number drawn from a uniform distribution on [0,1]. To generate the simulated FRET time traces and ensemble averaged Fourier spectra shown in **Fig. 5b-d**, we solve the above equations using the 4th order Runge-Kutta method with $f = 0.09\ \mathrm{hr}^{-1}$ (**Fig. 5b**), $f = 0.36\ \mathrm{hr}^{-1}$ (**Fig. 5c**), or $f = 0.27\ \mathrm{hr}^{-1}$ (**Fig. 5d**) to yield solutions $[\mathrm{MAPK}_f(t)]$.

To simulate resonance selectivity, we model two substrates, Substrate A and Substrate B, as driven two component linear oscillators with repressors $R_i$, as described above in **SD V**:

$$\begin{bmatrix} \dot{\mathrm{A}} \\ \dot{R}_\mathrm{A} \end{bmatrix} = \begin{bmatrix} -\beta_{\mathrm{A1}} & -\alpha_{\mathrm{A1}} \\ \alpha_{\mathrm{A2}} & -\beta_{\mathrm{A2}} \end{bmatrix} \begin{bmatrix} \mathrm{A} \\ R_\mathrm{A} \end{bmatrix} + \begin{bmatrix} \zeta_1 \\ \zeta_2 \end{bmatrix} [\mathrm{MAPK}_f(t)] \quad \text{(S8.2.1)}$$

$$\begin{bmatrix} \dot{\mathrm{B}} \\ \dot{R}_\mathrm{B} \end{bmatrix} = \begin{bmatrix} -\beta_{\mathrm{B1}} & -\alpha_{\mathrm{B1}} \\ \alpha_{\mathrm{B2}} & -\beta_{\mathrm{B2}} \end{bmatrix} \begin{bmatrix} \mathrm{B} \\ R_\mathrm{B} \end{bmatrix} + \begin{bmatrix} \zeta_1 \\ \zeta_2 \end{bmatrix} [\mathrm{MAPK}_f(t)]. \quad \text{(S8.2.2)}$$

We set $\zeta_2 = 0$ and use

$$\alpha_{\mathrm{A1}} = 1.6\ \mathrm{hr}^{-1}$$
$$\alpha_{\mathrm{A2}} = 0.8\ \mathrm{hr}^{-1}$$
$$\beta_{\mathrm{A1}} = \beta_{\mathrm{A2}} = 0.004\ \mathrm{hr}^{-1}$$
$$f_{\mathrm{A0}} = \frac{\sqrt{\alpha_{\mathrm{A1}}\alpha_{\mathrm{A2}} + \beta_{\mathrm{A1}}\beta_{\mathrm{A2}}}}{2\pi} = 0.18\ \mathrm{hr}^{-1}$$
$$Q_\mathrm{A} = \frac{\sqrt{\alpha_{\mathrm{A1}}\alpha_{\mathrm{A2}} + \beta_{\mathrm{A1}}\beta_{\mathrm{A2}}}}{\beta_{\mathrm{A1}} + \beta_{\mathrm{A2}}} = 141$$

and

$$\alpha_{\mathrm{B1}} = 3.2\ \mathrm{hr}^{-1}$$
$$\alpha_{\mathrm{B2}} = 1.6\ \mathrm{hr}^{-1}$$
$$\beta_{\mathrm{B1}} = \beta_{\mathrm{B2}} = 0.004\ \mathrm{hr}^{-1}$$
$$f_{\mathrm{B0}} = \frac{\sqrt{\alpha_{\mathrm{B1}}\alpha_{\mathrm{B2}} + \beta_{\mathrm{B1}}\beta_{\mathrm{B2}}}}{2\pi} = 0.36\ \mathrm{hr}^{-1}$$
$$Q_{\mathrm{B}} = \frac{\sqrt{\alpha_{\mathrm{B1}}\alpha_{\mathrm{B2}} + \beta_{\mathrm{B1}}\beta_{\mathrm{B2}}}}{\beta_{\mathrm{B1}} + \beta_{\mathrm{B2}}} = 224$$

### SD IX. Resonance and Impedance in Alternating Current (AC) Circuits

To draw comparisons between the MAPK cascade and electronic telecommunications systems, we first review the frequency-dependent behavior of simple AC circuits.

Series resistor-inductor-capacitor (RLC) electronic circuits, when driven by a sinusoidal voltage $V(t)$ at angular frequency $\omega$, generate an alternating current, $I(t) = \dot{Q}(t)$, described by

$$\ddot{Q}(t) + \frac{R}{L}\dot{Q}(t) + \frac{1}{LC}Q(t) = \frac{1}{L}V(t), \tag{S9.1}$$

where $R$ is the resistance, $L$ the inductance, and $C$ the capacitance. With the identifications

$$x = Q(t) \tag{S9.2.1}$$
$$\omega_0 = (LC)^{-1/2} \tag{S9.2.2}$$
$$\gamma = R/L \tag{S9.2.3}$$
$$F(t) = \frac{1}{L}V(t), \tag{S9.2.4}$$

eq. S9.1 becomes

$$\ddot{x} + \gamma\dot{x} + \omega_0^2 x = F(t), \tag{S9.3}$$

the general equation of a driven damped linear harmonic oscillator, eq. S4.1. Differentiating again yields an equation for the current

$$\ddot{I} + \gamma\dot{I} + \omega_0^2 I = -i\omega F(t), \tag{S9.4}$$

such that the solution can be expressed as

$$I(t) = -i\omega\chi(\omega)\frac{1}{L}V(t). \tag{S9.5}$$

Note the similarity to eq. S6.9. Therefore,

$$I(t) = Y(\omega)V(t) \tag{S9.6}$$

$$\equiv \frac{V(t)}{Z(\omega)}. \quad \text{(S9.7)}$$

This defines the complex frequency-dependent impedance of the circuit, $Z(\omega)$,

$$Z(\omega) = R + iX(\omega) = e^{i\phi_Z}\sqrt{R^2 + X^2(\omega)}, \quad \text{(S9.8)}$$

where $R$ is the resistance of the circuit and $X(\omega)$ the reactance, which depends on $L$ and $C$. The impedance is the inverse of the admittance, or the transfer function of the circuit,

$$Z(\omega) = Y^{-1}(\omega). \quad \text{(S9.9)}$$

The current in the driven circuit is given in terms of the driving voltage as in eq. S4.5

$$I(t) = Y(\omega)V(t) = \frac{V(t)}{Z(\omega)} = \frac{V(t)e^{-i\phi_Z}}{\sqrt{R^2 + X^2(\omega)}} = \frac{V(t)e^{-i\phi_Z}}{\sqrt{\gamma^2\omega^2 + (\omega_0^2 - \omega^2)^2}}, \quad \text{(S9.10)}$$

*i.e.*, the AC equivalent of Ohm's Law, with $\phi_Z$ the phase shift between the current and voltage.

### SD X. The MAPK Cascade as Oscillating Phosphate Current Amplifier and Distribution Network

The oscillating components of the MAPK cascade carry negative electrically charged phosphate residues being added and removed from components by enzymatic chemical reactions, constituting a literal electrical oscillating current. In this section we apply principles of AC circuit behavior to the MAPK cascade to describe how the MAPK systems amplifies and distributes an AC current of phosphates to elements in the cascade to control gene expression.

Because of the close correspondence between eq. S6.9 and eq. S9.5, we identify $x_j(t)$ and $z(t)$ as phosphate currents. The $\mu_j(t)$ in eqs. S5.12, S5.17, and S5.18 represent chemical potentials induced by the upstream $z(t)$ current to drive the $x_j(t)$ currents,

$$\mu_j(t) = \left(\beta_k\zeta_j \mp \alpha_k\zeta_k\right)z + \zeta_j\dot{z}. \quad \text{(S10.1)}$$

The first term represents DC coupling through Ohm's law,

$$V_1(t) = RI, \quad \text{(S10.3)}$$

which, as described above (see **SD V**), sets the average activation level of the TF.

The second term represents coupling by mutual inductance,

$$V_1(t) = M\dot{I}_2, \quad \text{(S10.2)}$$

or how current changes in one element generate an AC voltage in an inductively coupled second element, resulting in oscillations around the average activation level. The net chemical potential is determined as the superposition of all input potentials (**Fig. 5g**).

Consequently, each oscillating component in the MAPK cascade can behave exactly as an electrical AC circuit element with complex impedance equal to the reciprocal of its admittance/transfer function, $Z_j(\omega) = Y^{-1}(\omega)$.

Each component is coupled to other components, such that the current in that element is

$$x_j(t) = \sum_k Y_{jk}(\omega) z_k(t) = \sum_k \frac{z_k(t)}{Z_{jk}(\omega)} \tag{S10.4}$$

with the complex impedance $Z_k(\omega)$ encapsulating the resonant response of the element

$$Z_k(\omega) = e^{i\phi_{Z_k}} \sqrt{\gamma^2\omega^2 + \left(\omega_{0,k}^2 - \omega^2\right)^2} \tag{S10.5}$$

$$\phi_{Z_k} = \tan^{-1}\left(\frac{\gamma\omega}{\omega_{0,k}^2 - \omega^2}\right). \tag{S10.6}$$

A component of the MAPK cascade interacting with a single upstream driver will carry oscillating phosphate current given by,

$$x_j(t) = Y_j(\omega) z(t) = \frac{z(t)}{Z_j(\omega)}, \tag{S10.7}$$

which corresponds to a gain of

$$\text{Gain} = \frac{|x_j(t)|}{|z(t)|} = \frac{1}{|Z_j(\omega)|}. \tag{S10.8}$$

As we find experimentally $\gamma \ll 1$, $Q \gg 1$, the gain will only be significant near resonance ($\omega \approx \omega_0$), yielding

$$\text{Gain} = \frac{\zeta_j}{\gamma_j}. \tag{S10.9}$$

Gain greater than one, corresponding to the component acting as a phosphate current amplifier, requires $\zeta_j > \gamma_j$, that is, the catalytic rate of the driving enzyme phosphorylating the targets must be greater than the sum of the spontaneous dephosphorylation rates of the targets.

For multiple components in series, the driving current $z(t)$ in eq. S10.7 will itself have been generated in response to a further upstream component. Thus, $N$ components in series generate a net gain

$$\text{Net gain} = \prod_{j=1}^{N} Z_j^{-1}(\omega) \approx \prod_{j=1}^{N} \frac{\zeta_j}{\gamma_j}, \tag{S10.10}$$

That is, phosphate current paths through the cascade that are near resonance (*i.e.*, low net impedance) result in large amplification of the phosphate current along that path. Conversely, with high $Q$ values, the impedance of pathways composed of elements off-resonance for a given signal will experience high resistance/impedance and low amplification. This is illustrated in **Fig. 5g** as the magnitude of the current along a resonant path increasing in thickness along the cascade.

### SD XI. Frequency Shift Keying (FSK) and Orthogonal Frequency-Division Multiplexing (OFDM)

An encoding scheme commonly used in digital telecommunications systems is Frequency Shift Keying (FSK), where switching the state of a digital bit from 0 to 1 is communicated by changing the frequency of a carrier wave from $f_0$ to $f_1$ (or *vice versa*). Our results suggest that p38's information encoding scheme is similar, where the presence or absence of stimuli is communicated by switching p38 from its resting state to an active state with oscillatory frequency $f_i$. Moreover, specific types of stimuli experienced are encoded in which frequencies appear in p38's spectrum. This scheme is a form of multiplexed FSK, where the state of multiple bits is encoded by the appearance of specific frequencies in the overall spectrum of the carrier wave. This correspondence is not exact; p38's encoding is analog rather than digital, *i.e.*, the magnitude of the stimulus is also encoded in the amplitude of p38's response (**Fig. 1e,f**). However, the comparison is still instructive.

Each frequency in the carrier wave is detected by a receiver. Here, this would correspond to p38 substrates that oscillate at a specific natural frequency due to their interactions with other elements in the cell (see **SD V**). To maximize the fidelity of information transmission, each receiver should ideally respond to only one input frequency. This is accomplished electronically by using orthogonal frequencies, corresponding to the encoding scheme referred to as Orthogonal Frequency-Division Multiplexing (OFDM).

Consider two signal waves, $\Psi_j(t)$ and $\Psi_k(t)$, with corresponding frequencies $f_j$ and $f_k$. These signals are orthogonal if their time-averaged correlation is non-zero only if the frequencies are identical:

$$\langle \Psi_j(t), \Psi_k(t) \rangle \equiv \frac{1}{T_S} \int_0^{T_S} \Psi_j(t) \cdot \Psi_k^*(t) dt = \delta_{jk}, \tag{S11.1}$$

where the asterisk indicates the complex conjugate, $\delta_{jk}$ is the Kronecker delta,

$$\delta_{jk} = \begin{cases} 0 & j \neq k \\ 1 & j = k \end{cases}, \tag{S11.2}$$

and $T_s$ is the symbol period, *i.e.*, the time required to transmit the state of a bit. Since the time-averaged correlation between two orthogonal signal waves with different frequencies is zero, use of orthogonal frequencies results in the minimum possible crosstalk between receivers.

Note that orthogonality is satisfied if the frequencies used are harmonics, which are evenly spaced integer multiples of a fundamental frequency. For example,

$$\int_0^{\frac{2\pi}{\omega_0}} \cos(m\omega_0 t)\cdot\cos(n\omega_0 t)\,dt = \frac{\pi}{\omega_0}\delta_{m,n}, \tag{S11.3}$$

with *m* and *n* integers. Hence the harmonic frequencies employed by p38 are orthogonal and would provide the minimum possible crosstalk between TFs responding at different frequencies in an OFDM encoding scheme.

The symbol period in an OFDM scheme is the inverse of the frequency spacing. In the case of p38 harmonics, the frequency spacing is the fundamental frequency $f_0 = 0.09\ \mathrm{hr}^{-1}$, such that $T_s = f_0^{-1} \cong 11$ hours. For this reason, we collected RNA samples for sequencing after 48 hours, or approximately four symbol periods, to ensure that sufficient time was allowed for signals to be fully communicated and the response initiated.

OFDM receivers are typically constructed to act as "correlator" circuits that perform the integral S11.1 to discriminate between input frequencies. The Michaelis-Menten description of the rate of product formation by an enzyme, eq. S5.2, provides the form required to act as a correlator receiver. In the regime $x_j' \ll K_j$ (with $K_j$ the binding constant of unphosphorylated substrate $x_j'$ with enzyme *z*), eq. S5.4, the driven two-component oscillator model eq. S5.8 would appear as

$$\frac{dx_1}{dt} = -\alpha_2 x_2 - \beta_1 x_1 + \zeta_1' x_1' z \tag{S11.4.1}$$

$$\frac{dx_2}{dt} = \alpha_1 x_1 - \beta_2 x_2 + \zeta_2' x_2' z. \tag{S11.4.2}$$

If all components oscillate, then taking the time average of eqs. S11.4 results in the last term acting as a correlator receiver, eq. S11.1. Hence the time-averaged rate of change of each component will only be non-zero if the frequency of the driver matches the natural frequency of the TF substrate.
